# Cloud-based DIA data analysis module for signal refinement improves accuracy and throughput of large datasets

**DOI:** 10.1101/2021.07.14.452243

**Authors:** Karen E. Christianson, Jacob. D. Jaffe, Steven A. Carr, Alvaro Sebastian Vaca Jacome

**Affiliations:** Broad Institute of MIT and Harvard, Cambridge, MA, USA; Inzen Therapeutics, Cambridge, MA, USA

## Abstract

Data-independent acquisition (DIA) is a powerful mass spectrometry method that promises higher coverage, reproducibility, and throughput than traditional quantitative proteomics approaches. However, the complexity of DIA data caused by fragmentation of co-isolating peptides presents significant challenges for confident assignment of identity and quantity, information that is essential for deriving meaningful biological insight from the data. To overcome this problem, we previously developed Avant-garde, a tool for automated signal refinement of DIA and other targeted mass spectrometry data. AvG is designed to work alongside existing tools for peptide detection to address the reliability and quantitative suitability of signals extracted for the identified peptides. While its use is straightforward and offers efficient refinement for small datasets, the execution of AvG for large DIA datasets is time-consuming, especially if run with limited computational resources. To overcome these limitations, we present here an improved, cloud-based implementation of the AvG algorithm deployed on Terra, a user-friendly cloud-based platform for large-scale data analysis and sharing, as an accessible and standardized resource to the wider community.

## Main Text

Data-independent acquisition (DIA) is a powerful mass spectrometry method that promises higher coverage, reproducibility, and throughput than traditional quantitative proteomics approaches ^1–3^. In DIA mode, the instrument fragments and analyzes all co-isolating peptides as it cycles through windows of a given m/z range, thereby generating time-resolved fragment ion spectra for all peptides above the instrument’s limit of detection ^2,4–6^. In principle, DIA combines the sensitivity and throughput of targeted methods with the higher proteome coverage of data-dependent acquisition. However, the complexity of the acquired DIA data, caused by fragmentation of co-isolating peptides and use of a limited number of fragment ions for identification (sometimes as few as 4), presents significant challenges for confident assignment of identity and quantity, information that is essential for deriving biological insight from the data^1,3,6,7^. A widely accepted method to address this problem in DIA data analysis is targeted extraction of fragment ion chromatograms for peptides based on prior observation in a curated spectral library obtained by deep-scale data-dependent LC-MS/MS analysis ^1,3,8–10^.

We previously developed Avant-garde (AvG) ^11^, a tool for automated signal refinement of DIA and other targeted mass spectrometry data. AvG is designed to work alongside existing tools for peptide detection ^8,12–15^ to address the reliability and quantitative suitability of signals extracted for the identified peptides. Following targeted chromatogram extraction of identified peptides, AvG refines chromatographic peak traces by removing interfering transitions, adjusting integration boundaries, and scoring peaks to control false discovery rate, thereby increasing confidence in the resultant quantitation^11^. AvG is not limited to DIA data and can be used to curate any data that produces fragment-ion chromatograms at the MS2 level, such as Parallel Reaction Monitoring (PRM).

AvG was initially designed for seamless integration with Skyline ^16^, a vendor-independent and user-friendly software for visualization and analysis of targeted proteomics experiments. AvG is available for download as an external tool in the Skyline Tool Store and can be used directly within a Skyline document. While the implementation of AvG in Skyline is straightforward and offers efficient refinement for small datasets (i.e. PRM data or DIA datasets with limited samples and peptide targets), the execution of AvG for large DIA datasets (i.e. thousands of peptide targets) is time-consuming, especially if run with limited computational resources. Motivated by the need to analyze large DIA datasets, we initially implemented the AvG algorithm locally on a parallel computing platform as described in the original publication^11^. However, such large compute clusters may not be commonly accessible and their use requires advanced technical knowledge.

To overcome these limitations, we present here an improved, cloud-based implementation of the AvG algorithm deployed on Terra (https://app.terra.bio), a user-friendly cloud-based platform for large-scale data analysis and sharing, as an accessible and standardized resource to the wider community. The new AvG workflow is optimized for large DIA datasets and uses the Terra workflow management to fully automate all processing steps so that no user interaction is required after uploading the data to be analyzed. Terra provides workflow management and facilitates the assembly of long-running pipelines comprising a large number of tasks that typically require days or even weeks to complete. Each step of the AvG algorithm was implemented as a task, and the workflow represents the series of tasks (a pipeline) where outputs of a given task are the inputs of the next task. The workflow is written in the Workflow Description Language (WDL) and executes dockerized R and Python tools for pre-processing, running the AvG algorithm, and post-processing, all of which are available in a public docker image (Figure 1 and Supplementary Figure 1).

**Figure 1:**
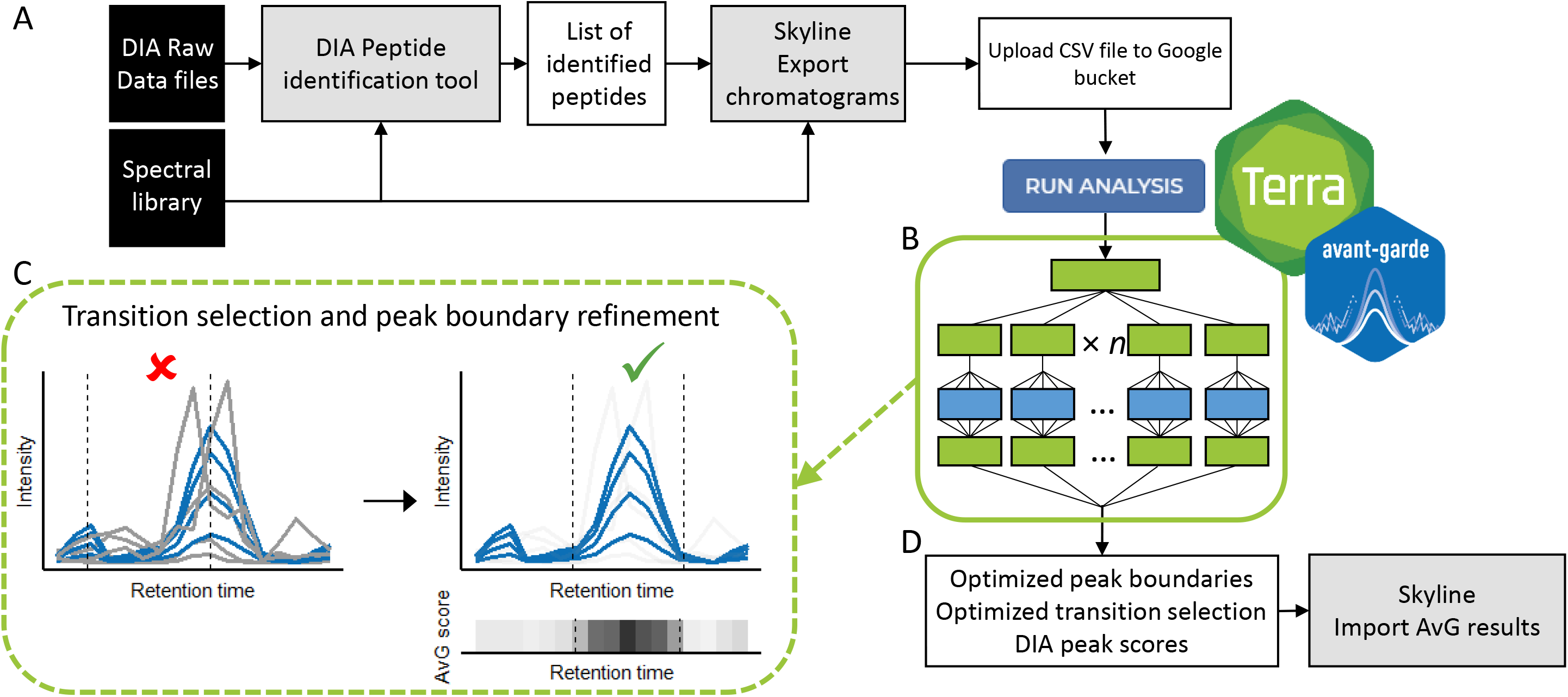
Overview of Avant-garde Terra framework. Avant-garde works alongside DIA identification tools to address the quantitative suitability of signals extracted for the identified peptides. A. Skyline is used to extract chromatogram data of detected peptides. B. The workflow consists of 3 separate tasks represented by the solid green outline: pre-processing and formatting the input dataset, AvG’s genetic algorithm optimization, and post-processing to finalize results. For the pre-processing step, the user simply provides an input CSV file containing chromatogram signal information exported from Skyline. The data is transformed into the Apache Parquet file format for more efficient compression, indexing and partitioning of the data, allowing for more efficient parallelization and scalability of the AvG algorithm. The AvG algorithm task is first parallelized onto a user-defined number of virtual machines and further parallelized on each machine to use all available CPU cores for maximum computing efficiency. Finally, post-processing combines AvG results across all machines into final CSV reports. All steps within the green outline are fully automated and require no user interaction (Supplementary Figure 1). C. AvG curates transitions to reduce noise and remove interference using a genetic algorithm, assigning a final quality metric to the selected set. It also refines peak integration boundaries by calculating chromatographic subscores at each time point in the raw data, and combining them as a weighted product (AvG chromatographic score), where the maximum value of this score corresponds to the most likely retention time of the analyte. D. The output of AvG (suitable transitions, chromatographic boundaries and scoring metrics) is reimported to Skyline to produce curated quantitative data.

The AvG Terra workspace is a self-contained computational sandbox with everything you need to complete a project. This includes the AvG workflow, user-defined analysis parameters, and the results. Additionally, reproducible results can be ensured by Terra’s version control system for algorithms, parameters, and results. The cloud-based AvG workflow harnesses the power of cloud-based computing to provide reproducible, user-friendly, and efficient processing of large DIA datasets with greater accessibility than our previous workflow. Its distribution on Terra allows anyone, regardless of coding experience, to perform efficient AvG analysis for large DIA datasets without the need to write code, run code locally using unfriendly command-line tools, or access large-scale on-premises computing resources.

The new cloud-based implementation of AvG consists of the following components:

- A Terra workspace (https://app.terra.bio/#workspaces/lincs-phosphodia/Avant-garde_Production_v1_0) that includes the WDL workflow for running the AvG algorithm.
- A Github repository (https://github.com/broadinstitute/Avant-garde-Terra) with code, documentation, and description of the Terra workflow.
- A Github repository (https://github.com/SebVaca/avg_utils) for the python package containing the underlying functions required for the new AvG workflow.
- A tutorial (Suppplementary Information and https://github.com/broadinstitute/Avant-garde-Terra/wiki/Tutorial) with an accompanying Terra workspace (https://app.terra.bio/#workspaces/lincs-phosphodia/Avant-garde_Tutorial) illustrating the application of Avant-garde to a small spike-in peptide DIA dataset of 96 phosphopeptides x 15 samples^11^.

The new cloud-based AvG workflow on Terra was applied to three datasets. We first analyzed a peptide spike-in calibration curve dataset using both the Skyline External Tool version and the cloud-based Terra version of AvG. The relative quantification of the two datasets matched (Supplementary figure 2) demonstrating that the two implementations of AvG provide the same results for the relative quantification across a broad concentration range. We next applied the AvG Terra workflow to a triple-proteome sample consisting of four mixture samples of three complex proteomes (Human, E. Coli and Yeast) that was described in the original AvG paper^11^. The results of this evaluation show excellent accuracy, as the measured ratios matched very closely the known protein ratios (Supplementary Figure 3, Supplementary Figure 4 and supplementary Table 1). Furthermore, we were able to validate more than 34,000 precursors in the whole dataset (Supplementary Table 1 and 2) and obtained excellent reproducibility (Supplementary Figure 5 and Supplementary table 2). To further evaluate the confidence of the results, we searched the same Triple-species proteome samples with a Pyrococcus furiosus proteome spectral library, which provides a method for detecting random hits^17^. As expected, no proteins were identified searching the Pyrococcus furiosus library, and the score distribution of the Pyrococcus furiosus and the decoy peptides overlapped, demonstrating very low ’intrinsic’ false discovery rate (Supplementary figure 6 and supplementary Table 3).

The Skyline External tool and the new Terra implementation of the Avant-garde algorithm are complementary to each other and can serve different purposes. The Skyline External tool version of AvG is ideal for small datasets, especially PRM. The Terra workflow makes AvG refinement possible on large DIA datasets that would be time-consuming with the Skyline External tool version, and the scalability potential provided by Terra allows the user to customize the degree of parallelization in the workflow according to timing and cost restraints (supplementary Table 4). In the future, the AvG WDL workflow could be easily incorporated into a larger-scale DIA data analysis pipeline developed on Terra.

## Code availability

All source code required to run the Avant-garde workflow on Terra (WDL workflow, Dockerfile, required scripts) can be found on the Avant-garde-Terra Github page at https://github.com/broadinstitute/Avant-garde-Terra. The python package containing the underlying functions required for the workflow can be found on Github at https://github.com/SebVaca/avg_utils (the tar.gz version of this package is located in the src folder of the Avant-garde-Terra Github page). The docker image that hosts all analysis scripts is publicly available on DockerHub and can be pulled from “broadlincsproteomics/avant-garde:v1_0”. The Avant-garde algorithm source code can be found at https://github.com/SebVaca/Avant_garde. The Terra production workflow can be found and cloned for use at https://app.terra.bio/#workspaces/lincs-phosphodia/Avant-garde_Production_v1_0. All documentation is located on the Avant-garde-Terra Github Wiki page at https://github.com/broadinstitute/Avant-garde-Terra/wiki. A tutorial for use can be found at https://github.com/broadinstitute/Avant-garde-Terra/wiki/Tutorial.

## Data availability

The data that support the findings of this study are available from the corresponding author upon request.

## Acknowledgements

We thank Michael Kirby for his help creating the dockerfile incorporating Apache PyArrow. We thank Malvina Papanastasiou, Karsten Krug, and D. R. Mani for their feedback and comments on the manuscript. This work was funded by U54 HG008097 to J.D.J. This work was also supported in part by grants from the National Cancer Institute (NCI) Clinical Proteomic Tumor Analysis Consortium grants NIH/NCI U24-CA210986 and NIH/NCI U01 CA214125 to S.A.C. and by a grant from the Dr. Miriam and Sheldon G. Adelson Medical Research Foundation to S.A.C. and Namrata D. Udeshi, Associate Director of the Proteomics Platform at the Broad Institute of MIT and Harvard.

## Contributions

K.E.C., J.D.J., and A.S.V.J. conceived the study. K.E.C. and A.S.V.J designed and executed the software, implementation, testing and documentation. A.S.V.J. designed and performed experiments and collected the data. J.D.J. provided laboratory resources, provided input on the software and provided guidance on the experimental design. S.A.C. provided laboratory resources and guidance on the manuscript. K.E.C and A.S.V.J and S.A.C wrote the manuscript, with input and revisions from all authors.

## Corresponding authors

Correspondence to Alvaro Sebastian Vaca Jacome or Steven A. Carr.

---

**Supplementary Figure 1.**
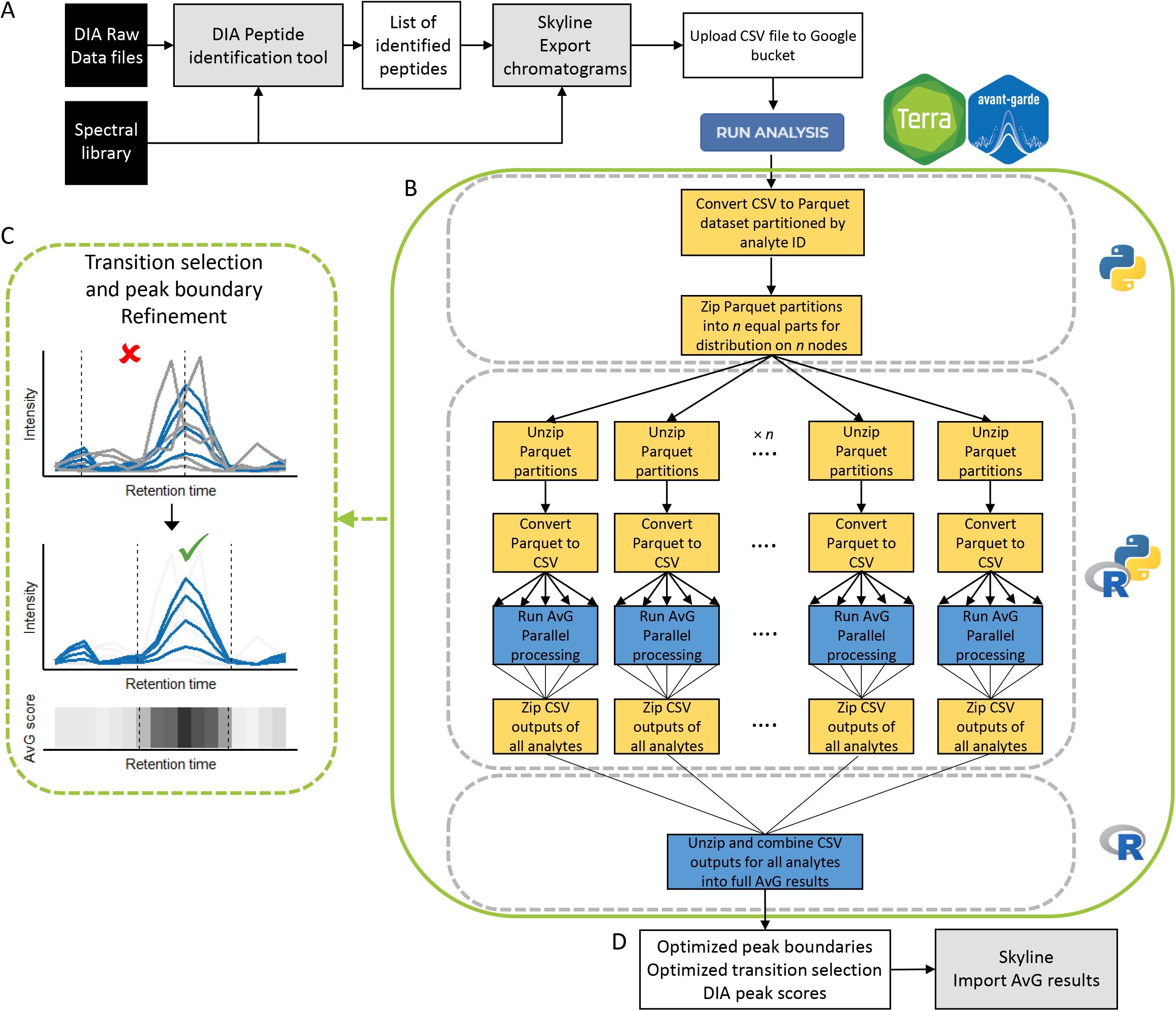

**Supplementary Figure 2.**
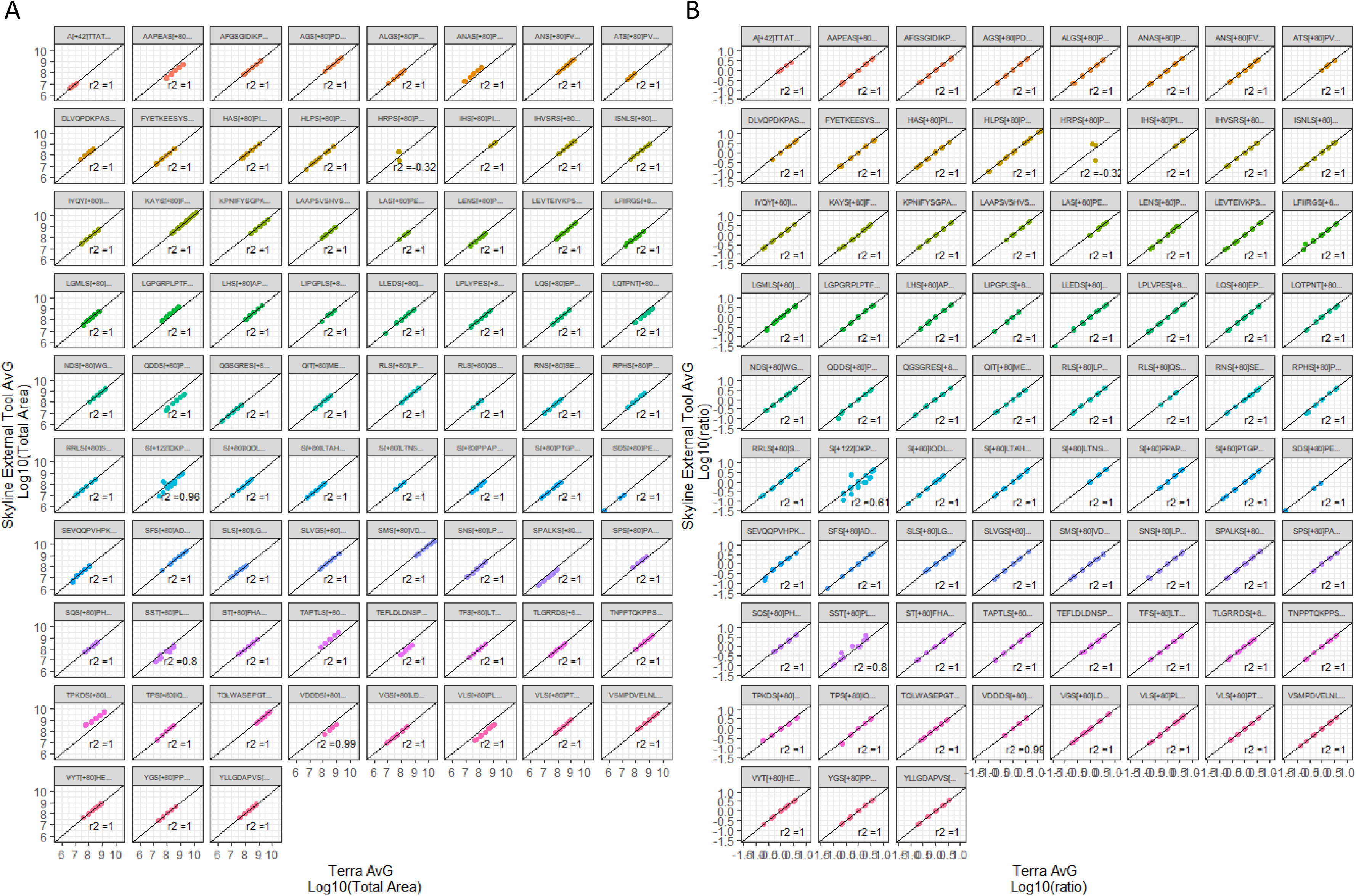

**Supplementary Figure 3.**
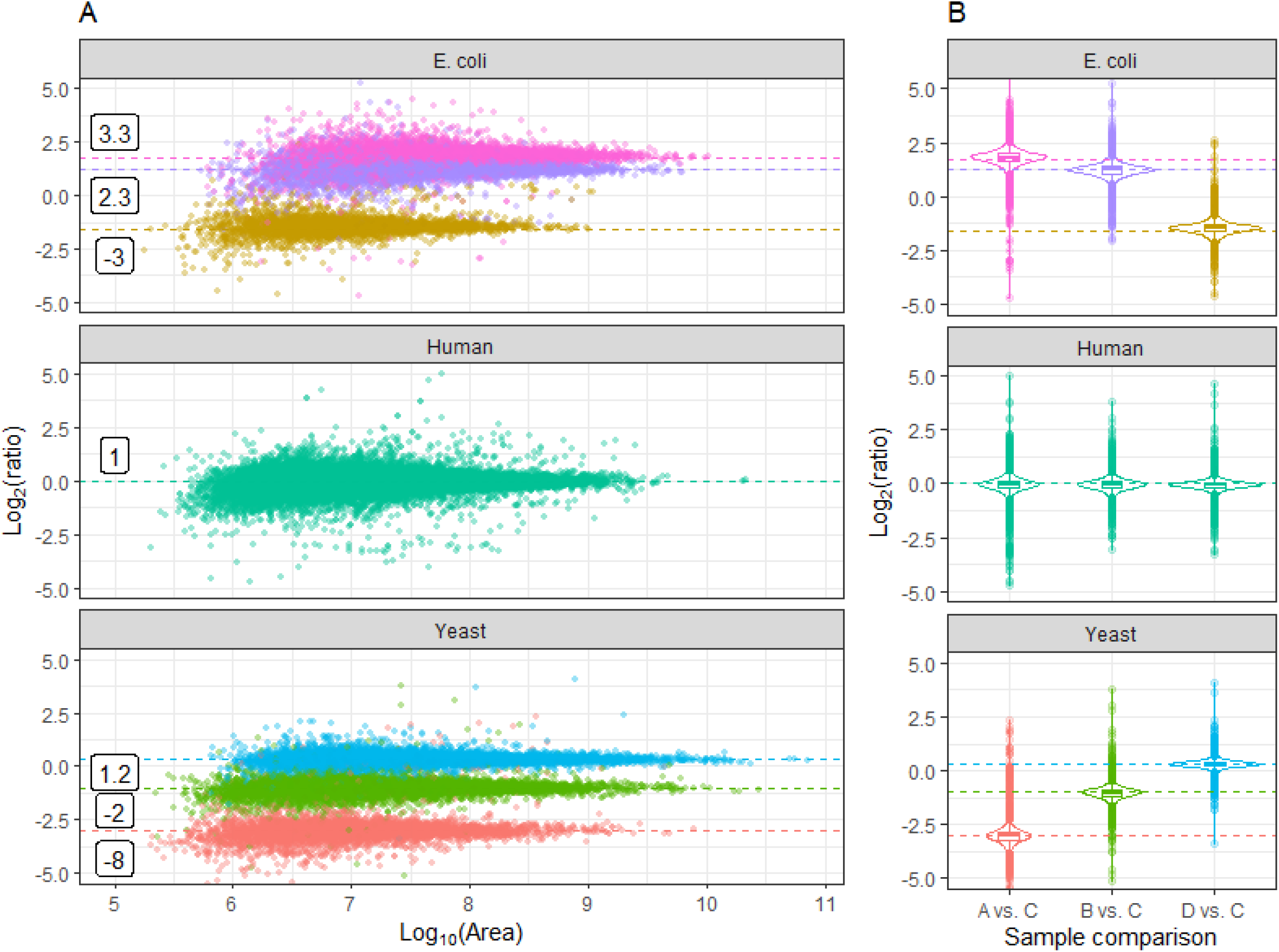

**Supplementary Figure 4.**
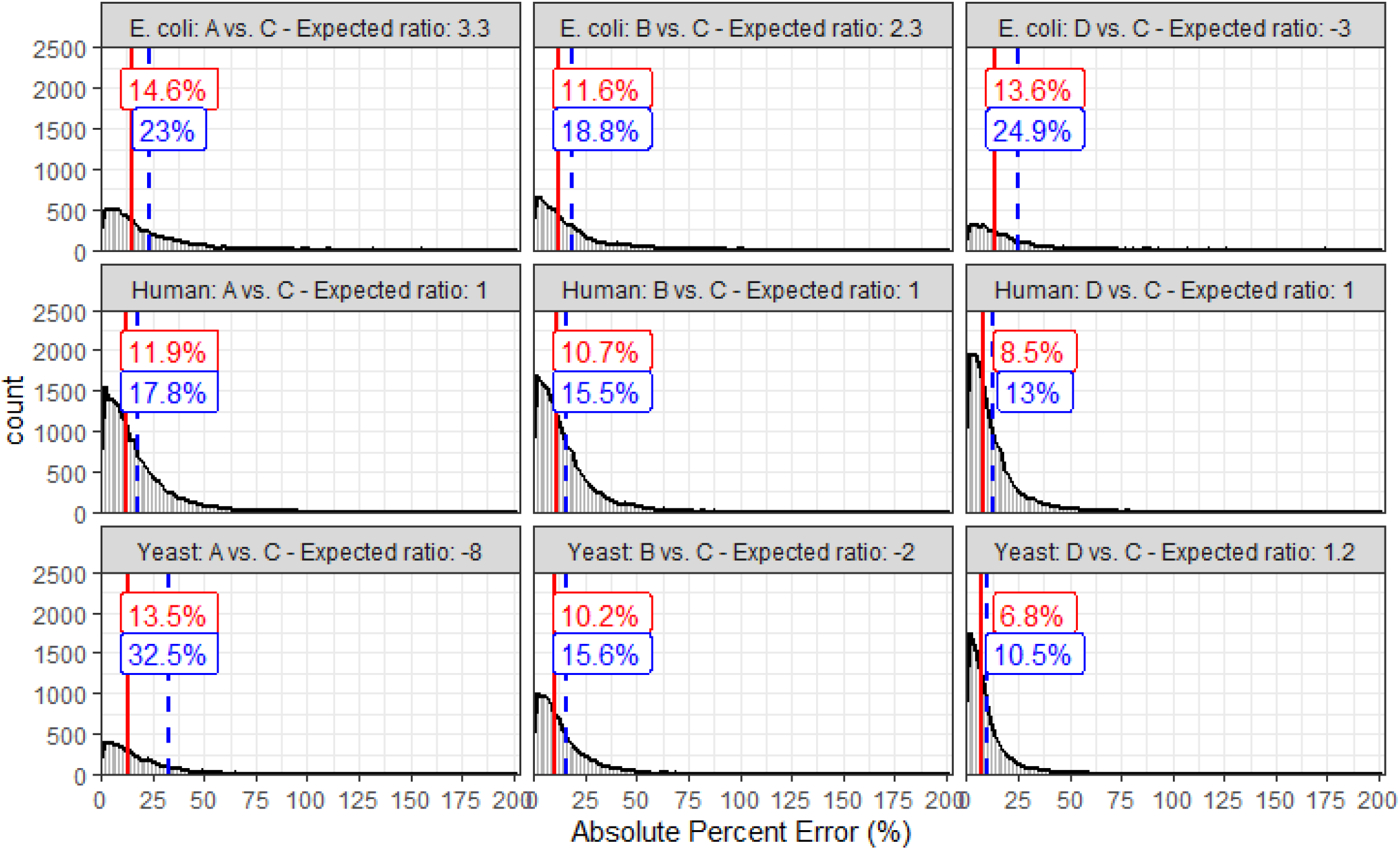

**Supplementary Figure 5.**
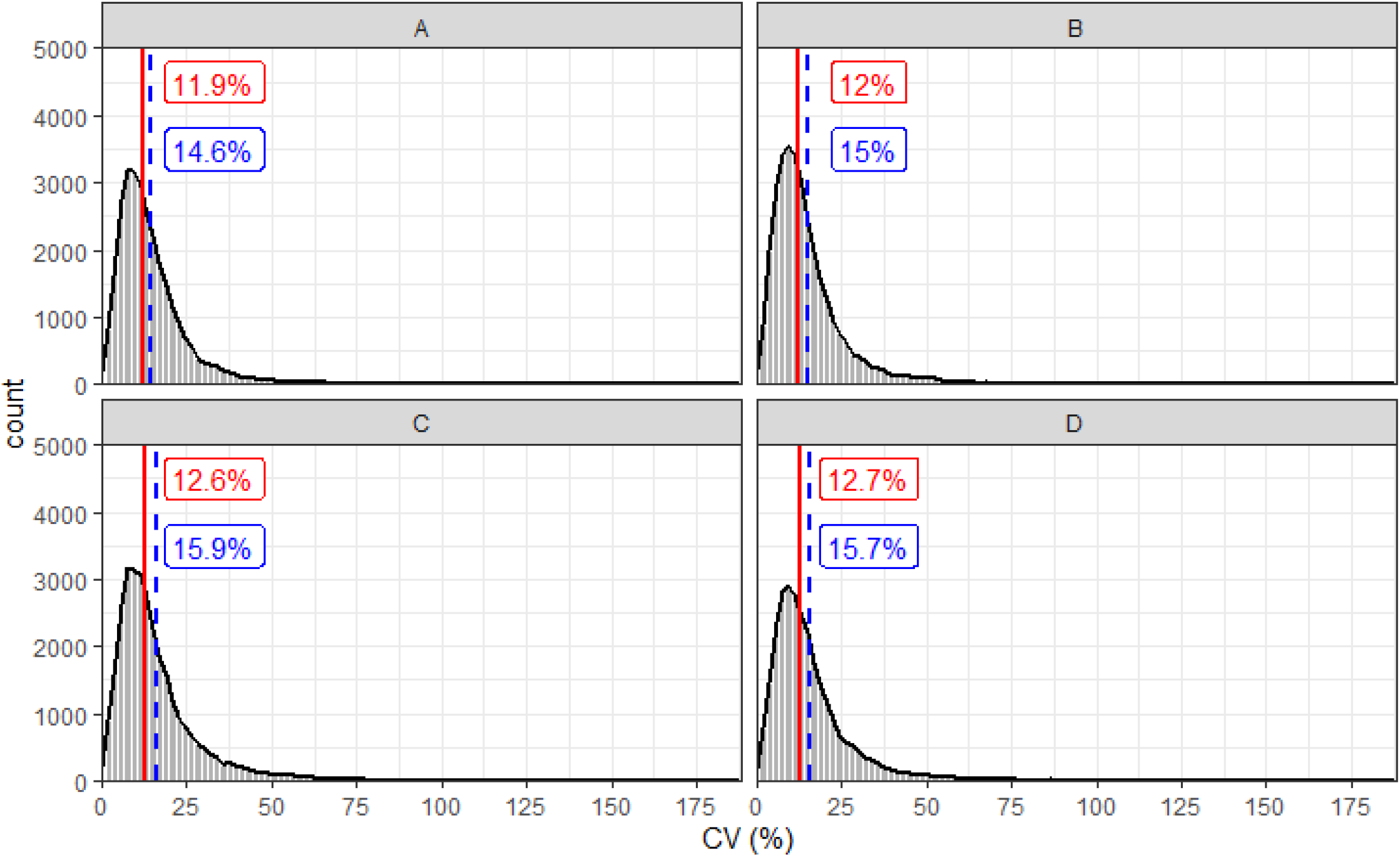

**Supplementary Figure 6.**
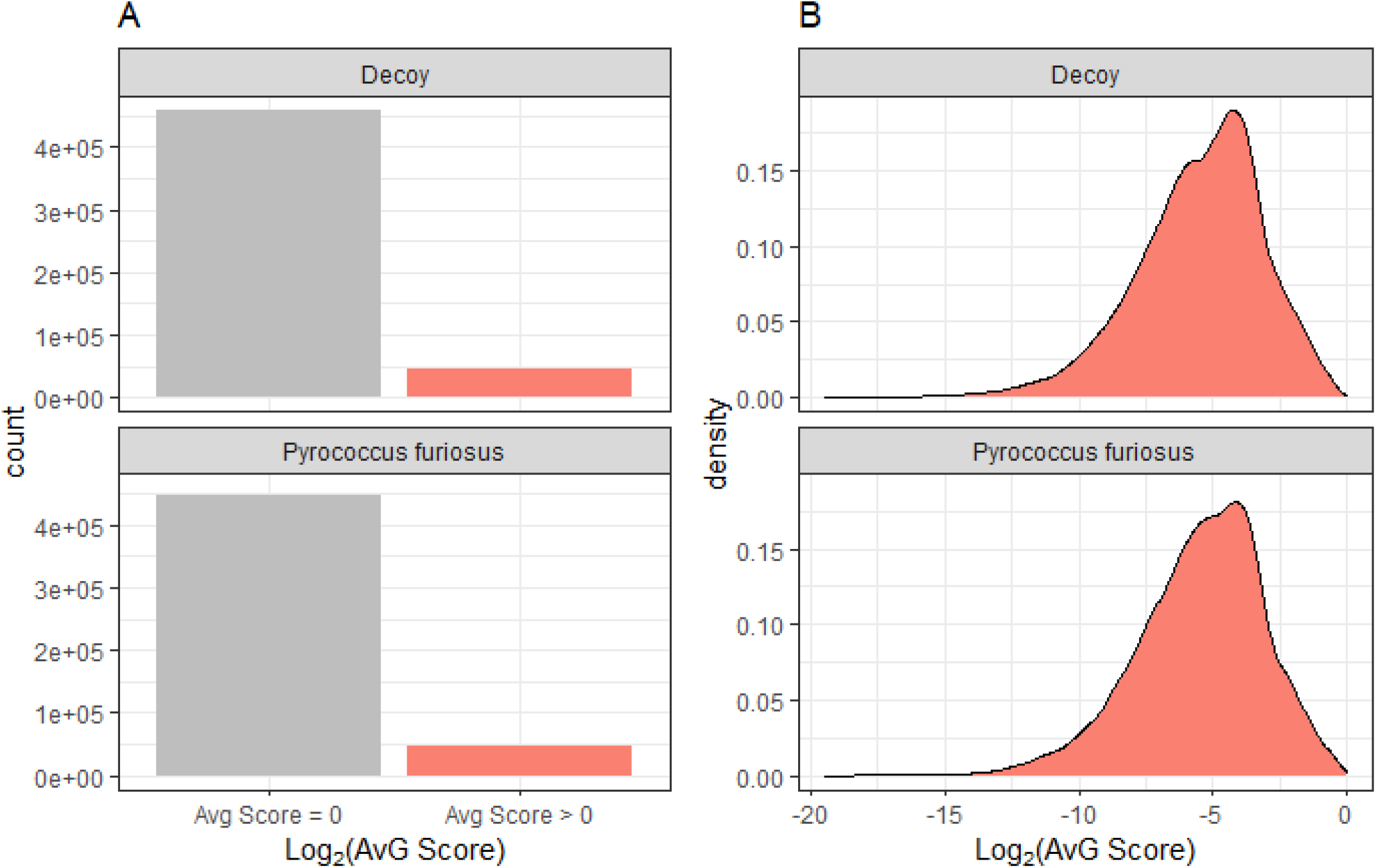

## References

1. Gillet, L. C. et al. Targeted data extraction of the MS/MS spectra generated by data-independent acquisition: a new concept for consistent and accurate proteome analysis. Mol. Cell. Proteomics 11, O111.016717 (2012).

2. Chapman, J. D., Goodlett, D. R. & Masselon, C. D. Multiplexed and data-independent tandem mass spectrometry for global proteome profiling. Mass Spectrom. Rev. 33, 452–470 (2014).

3. Röst, H. L., Aebersold, R. & Schubert, O. T. Automated SWATH Data Analysis Using Targeted Extraction of Ion Chromatograms. Methods Mol. Biol. 1550, 289–307 (2017).

4. Silva, J. C. et al. Quantitative proteomic analysis by accurate mass retention time pairs. Anal. Chem. 77, 2187–2200 (2005).

5. Silva, J. C. et al. Simultaneous qualitative and quantitative analysis of the Escherichia coli proteome: a sweet tale. Mol. Cell. Proteomics 5, 589–607 (2006).

6. Bilbao, A. et al. Processing strategies and software solutions for data-independent acquisition in mass spectrometry. Proteomics 15, 964–980 (2015).

7. Panchaud, A. et al. Precursor acquisition independent from ion count: how to dive deeper into the proteomics ocean. Anal. Chem. 81, 6481–6488 (2009).

8. Röst, H. L. et al. OpenSWATH enables automated, targeted analysis of data-independent acquisition MS data. Nat. Biotechnol. 32, 219–223 (2014).

9. Egertson, J. D., MacLean, B., Johnson, R., Xuan, Y. & MacCoss, M. J. Multiplexed peptide analysis using data-independent acquisition and Skyline. Nat. Protoc. 10, 887–903 (2015).

10. Schubert, O. T. et al. Building high-quality assay libraries for targeted analysis of SWATH MS data. Nat. Protoc. 10, 426–441 (2015).

11. Vaca Jacome, A. S. et al. Avant-garde: an automated data-driven DIA data curation tool. Nat. Methods 17, 1237–1244 (2020).

12. Reiter, L. et al. mProphet: automated data processing and statistical validation for large-scale SRM experiments. Nat. Methods 8, 430–435 (2011).

13. Tsou, C.-C. et al. DIA-Umpire: comprehensive computational framework for data-independent acquisition proteomics. Nat. Methods 12, 258–64, 7 p following 264 (2015).

14. Searle, B. C. et al. Chromatogram libraries improve peptide detection and quantification by data independent acquisition mass spectrometry. Nat. Commun. 9, 5128 (2018).

15. Peckner, R. et al. Specter: linear deconvolution for targeted analysis of data-independent acquisition mass spectrometry proteomics. Nat. Methods 15, 371–378 (2018).

16. MacLean, B. et al. Skyline: an open source document editor for creating and analyzing targeted proteomics experiments. Bioinformatics 26, 966–968 (2010).

17. Vaudel, M. et al. A complex standard for protein identification, designed by evolution. J. Proteome Res. 11, 5065–5071 (2012).

