## Supplementary Information for "Cloud-based DIA data analysis module for signal refinement improves accuracy and throughput of large datasets"

### Supplementary Figures

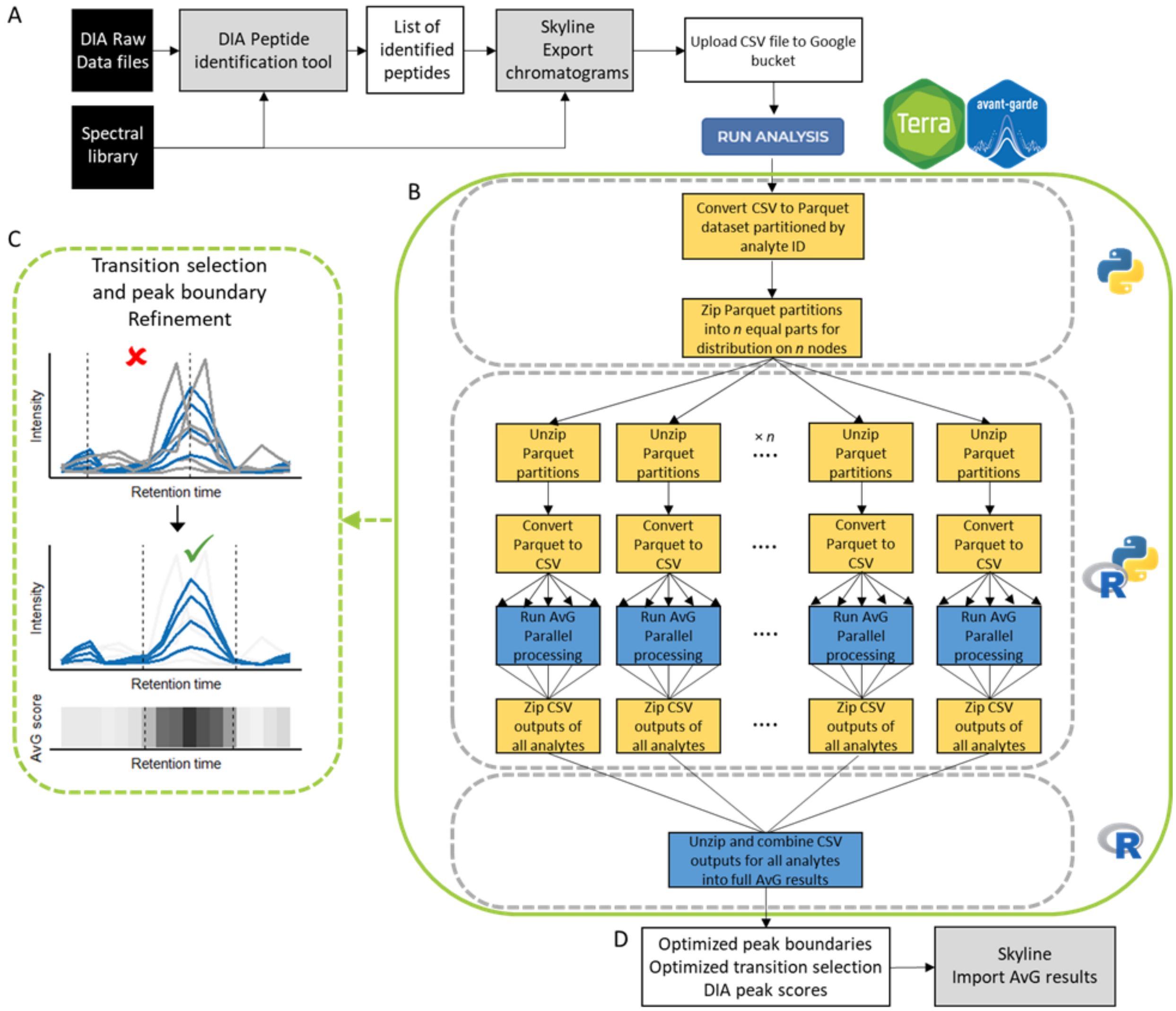


**Supplementary Figure 1. Overview of Avant-garde Terra framework.** Overview of the Avant-garde Terra workflow. Avant-garde is meant to work alongside DIA identification tools to address the quantitative suitability of signals extracted for the identified peptides. The input to the Terra workflow is a CSV file containing chromatogram signal information exported from Skyline. A. Skyline is used to extract chromatogram data of detected peptides. B. The workflow consists of 3 separate tasks outlined in dotted gray representing the pre-processing step, the AvG's genetic algorithm optimization, and the post-processing step. The user provides the input CSV which is first transformed into the Apache Parquet file format for a more efficient compression, indexing and partitioning of the data. This allows for more efficient parallelized computing making sure that only the necessary data is localized to the corresponding node in the second task. The second task runs the AvG algorithm in parallel over a user-defined number of machines, and further parallelization occurs on each machine to use all available CPUs for maximum computing efficiency. The final task combines AvG results from individual Parquet partitions into final CSV reports of AvG results. Each grey dashed-line box in this schematic describes one task in the workflow that has been dockerized; within these gray outlines, the yellow rectangular boxes represent steps executed in python and blue represent steps executed in R. All steps within the green box outline are fully automated and require no user interaction. C. AvG curates transitions to reduce noise and remove interference using a genetic algorithm, assigning a final quality metric to the selected set. It also refines peak integration boundaries by calculating chromatographic subscores at each time point in the raw data, and combining them as a weighted product (AvG chromatographic score), where the maximum value of this score corresponds to the most likely retention time of the analyte. D. The output of AvG (suitable transitions, chromatographic boundaries and scoring metrics) is reimported to Skyline to produce curated quantitative data.

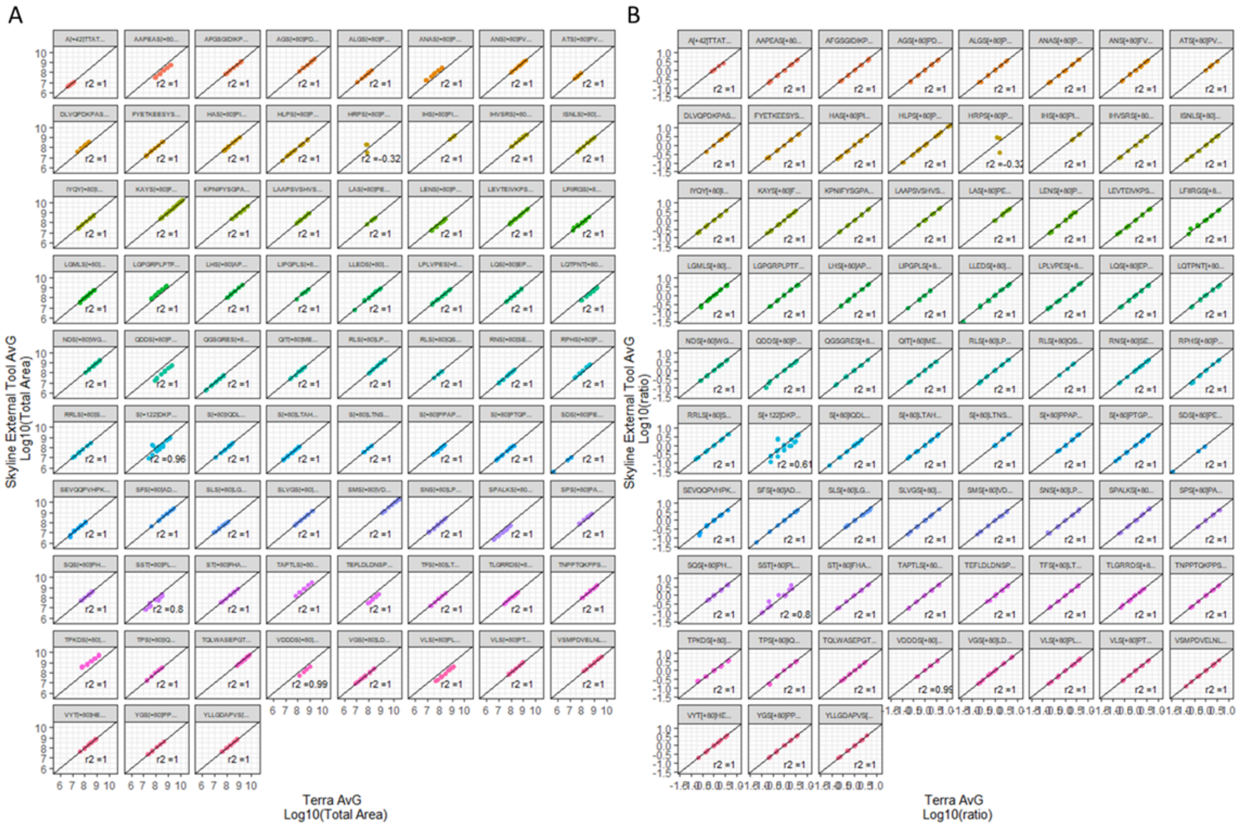


**Supplementary Figure 2: Comparison of results between the Skyline External Tool and the Terra implementation of Avant-garde.** Synthetic heavy isotope labeled peptides at known individual concentrations were spiked into a HEK293T digest to generate a five-point calibration curve. Each sample contained 1 µg HEK293T digest and 6.75, 13.5, 27, 54, or 108 ng total amount of heavy peptides. The five samples were analyzed in DIA mode in triplicates. In total, 15 runs were analyzed. (A). Each point represents the log10-transformed area obtained with the Skyline External Tool (Skyline External Tool  AvG, y-axis) and the Terra implementation of AvG (Terra AvG, x-axis). The black line represents the identity line (y = x). (B). Each point represents the ratio between the area to the median area in all 15 MS runs. The black line represents the identity line (y = x).

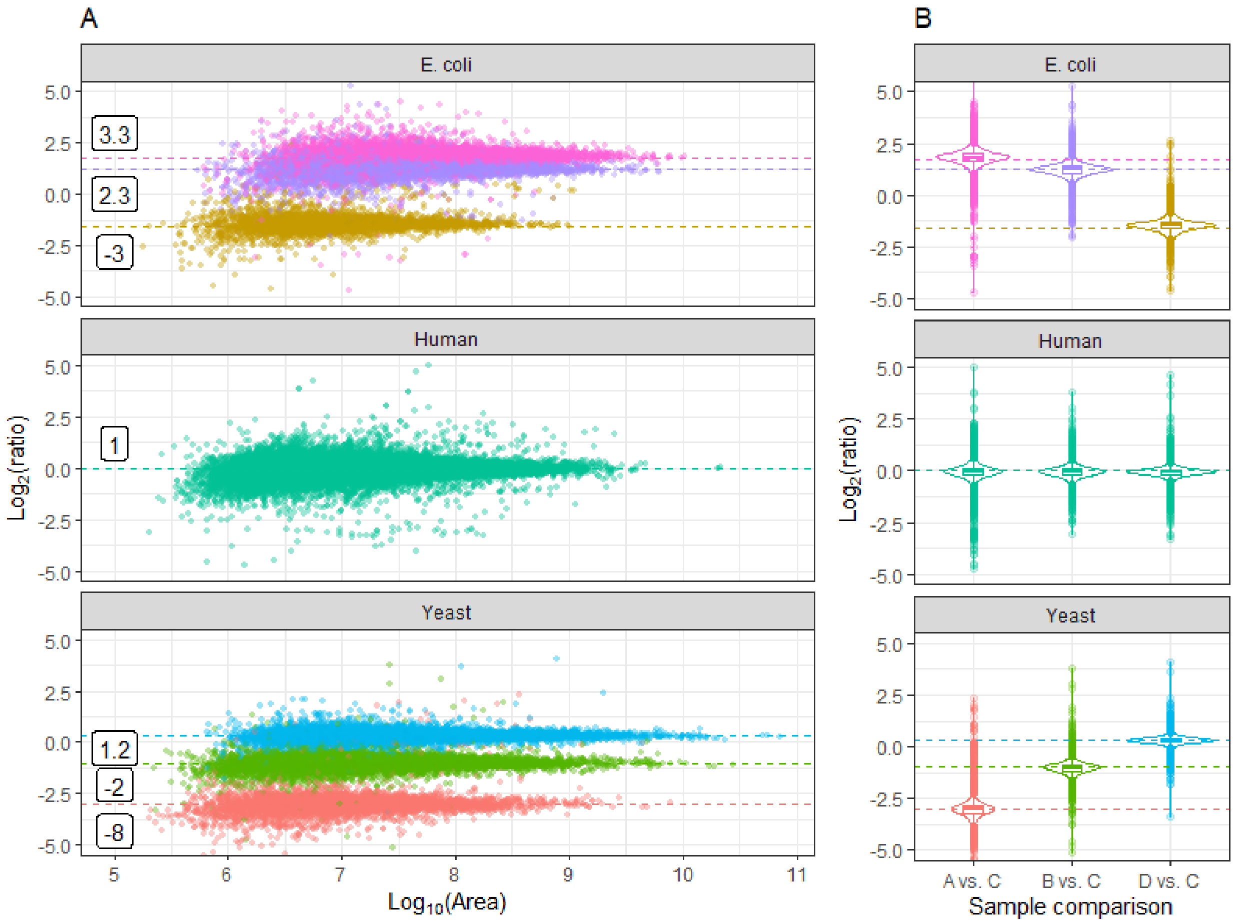


**Supplementary Figure 3: Relative quantification and accuracy evaluation of AvG using a triple-species benchmarking dataset.** (A). Distribution of ratios for the triple-proteome samples (n=4, analyzed in quadruplicate). The ratios were calculated using the mean area for each run (A, B, or D) divided by the mean area of sample C. Each dot represents a ratio calculated for a single peptide. The dashed horizontal lines represent the expected ratios, and the corresponding value of the fold change is in the white box. (B). The violin plot and boxplots show the distribution of the observed ratio of each pairwise combination (A vs. C, B vs. C and D vs. C). The box plot elements are: center line, median; box limits, upper and lower quartiles; whiskers, 1.5x interquartile range; points, outliers.

| Species | Comparison | Valid quantifiable ratio | Mean percent error (%) | Median percent error (%) |
|---|---|---|---|---|
| All | All | 34662 | 17.02 | 10.30 |
| E. coli | A vs. C | 6766 | 23.03 | 14.59 |
| E. coli | B vs. C | 6737 | 18.83 | 11.62 |
| E. coli | D vs. C | 3782 | 24.90 | 13.59 |
| E. coli | All | 6823 | 21.80 | 13.17 |
| Human | A vs. C | 15972 | 17.83 | 11.93 |
| Human | B vs. C | 16272 | 15.45 | 10.65 |
| Human | D vs. C | 15759 | 13.05 | 8.54 |
| Human | All | 17039 | 15.45 | 10.23 |
| Yeast | A vs. C | 4880 | 32.53 | 13.48 |
| Yeast | B vs. C | 9388 | 15.62 | 10.22 |
| Yeast | D vs. C | 10621 | 10.47 | 6.75 |
| Yeast | All | 10800 | 16.74 | 8.89 |

**Supplementary Table 1: Number of precursors and absolute value of the percent error in the triple-proteome sample analyzed by the Terra implementation of AvG.** The metrics are shown for each species, for each pairwise comparison and for the entire dataset (All).

| MS run | mean_CV | median_CV |
|---|---|---|
| A | 14.57 | 11.89 |
| B | 14.96 | 12.02 |
| C | 15.93 | 12.58 |
| D | 15.68 | 12.67 |
| All | 15.28 | 12.27 |

**Supplementary Table 2: Coefficient of variation in the triple-proteome sample analyzed by the Terra implementation of AvG.** The metrics are shown for each sample (A to D) and for the entire dataset (All).

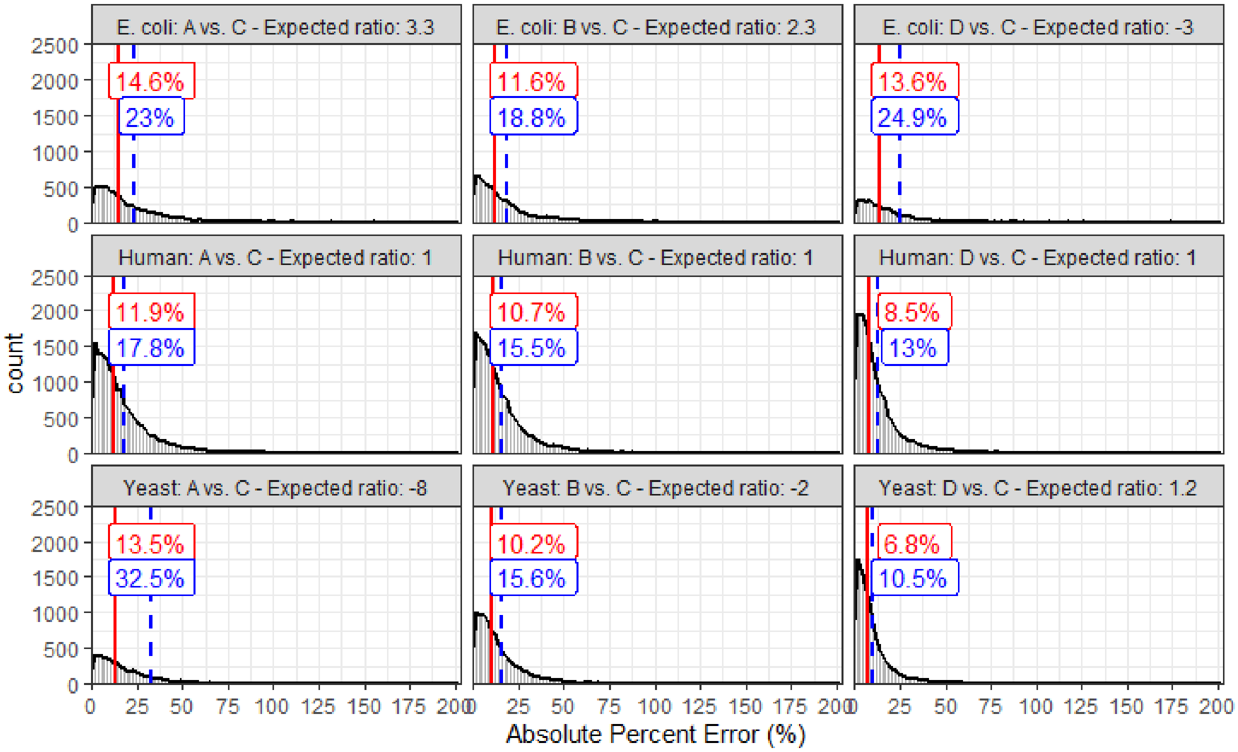


**Supplementary Figure 4: Evaluation of the accuracy of the relative quantification of the triple-proteome sample**. Distribution of absolute value of the percent error calculated using the ratio of a given peptide.The ratios were calculated using the mean area for each run (A,B or D) divided by the mean area of sample C. The percent error represents the deviation of the measured ratio to the expected ratio. The expected ratio is indicated on the title of each facet panel. Four replicates were analyzed for each peptide (n=4). The red full lines and the blue dashed lines show the median and the mean value respectively.

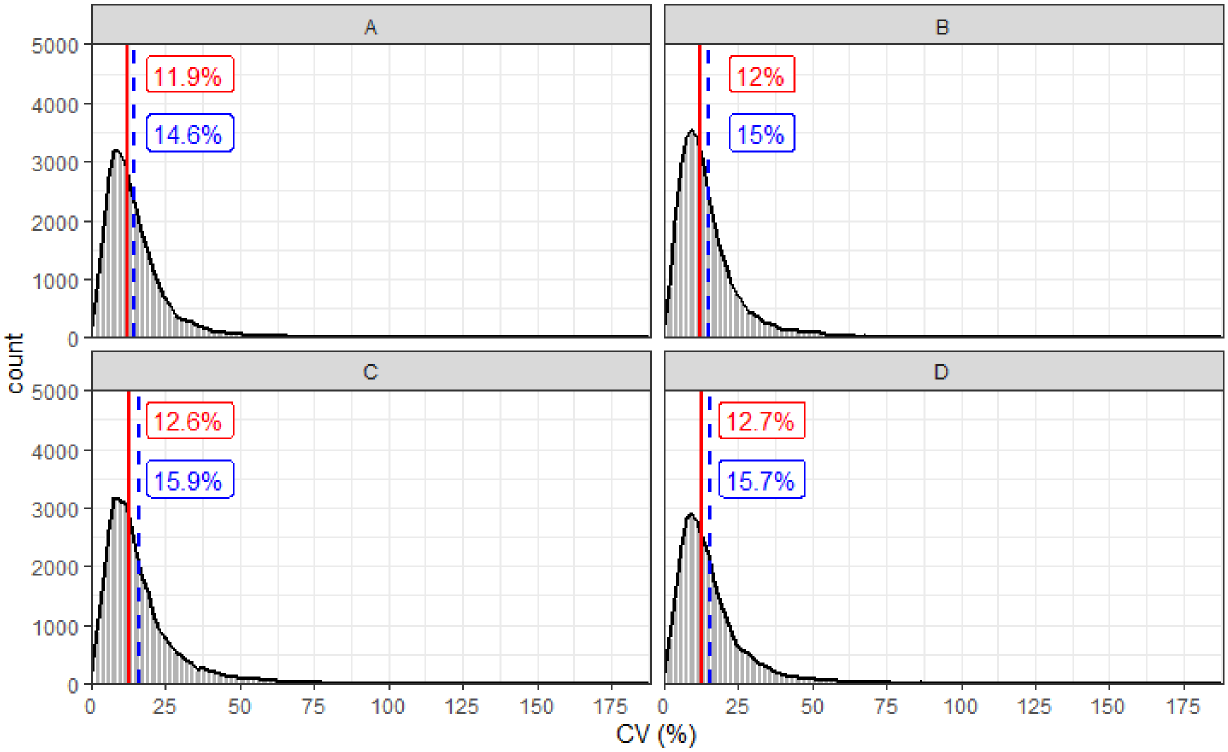


**Supplementary Figure 5: Evaluation of the precision of the relative quantification of the triple-proteome sample.** Distribution of coefficient of variation calculated using the areas in 4 replicates for each peptide. The red full lines and the blue dashed lines show the median and the mean value respectively.

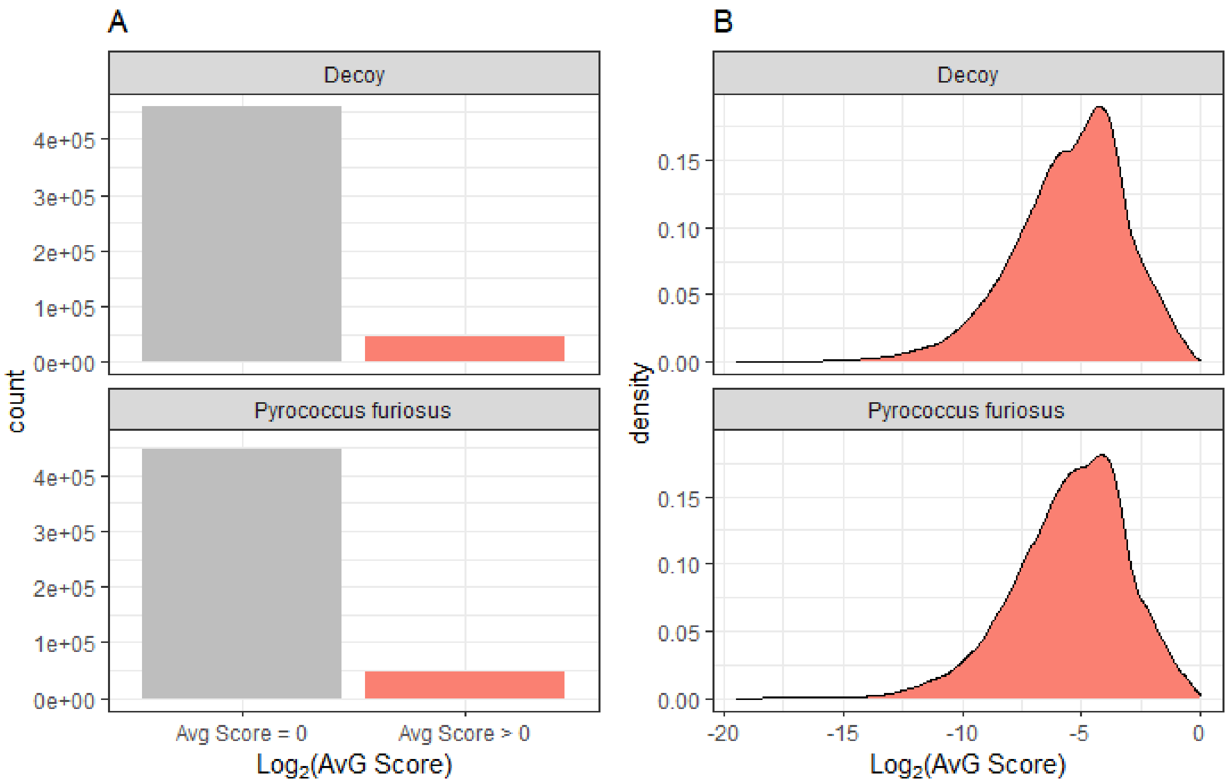


**Supplementary Figure 6: False discovery determination of the Terra implementation of AvG.** The triple-proteome dataset (E. Coli, yeast and human) was searched with a Pyrococcus furiosus proteome spectral library, which provides a method for detecting random hits. No Pyrococcus furiosus proteins were present in the sample. A. The distribution of the AvG scores for the decoy and Pyrococcus furiosus peptides is shown. B. A density plot for the non-zero AvG scores for the decoy and Pyrococcus furiosus peptides is shown.

| | Number of Pyrococcus furiosus DIA features | Number of Pyrococcus furiosus peptides |
|---|---|---|
| Total extracted | 497263 | 21074 |
| Validated (Filtered at q-value<0.01) | 15 | 10 |
| Validated (Filtered at PEP<0.05) | 0 | 0 |

**Supplementary table 3: Results of the false discovery determination of the Terra implementation of AvG.** We searched the triple-proteome dataset (E. Coli, yeast , human) with a Pyrococcus furiosus proteome strectral library, which provides a method for detecting random hits. No Pyrococcus furiosus proteins were present in the sample. The total number of DIA features extracted and validated and the corresponding number of peptides is shown. A DIA feature is defined as a peptide with a given charge state analyzed in a given MS run. The validation was done using Percolator v 3.0 and using a threshold of q-value<0.01 and posterior error probability<0.05 (PEP<0.05).

| Dataset name | Dataset size | Number of Peptides | Number of Precursors | Number of replicates | # virtual machines | # CPU per virtual machine | Run time | Run cost | cost per precursor ($) | Time per precursor (s) |
|---|---|---|---|---|---|---|---|---|---|---|
| Targeted | 189 MB | 96 | 96 | 15 | 1 | 30 | 0h 15m | $0.13 | $0.00135 | 9.38 |
| | | | | | 6 | 1 | 0h 19m | $0.07 | $0.00073 | 11.88 |
| | | | | | 6 | 2 | 0h 18m | $0.10 | $0.00104 | 11.25 |
| | | | | | 6 | 4 | 0h 15m | $0.11 | $0.00115 | 9.38 |
| | | | | | 6 | 6 | 0h 15m | $0.15 | $0.00156 | 9.38 |
| | | | | | 10 | 4 | 0h 13m | $0.16 | $0.00167 | 8.13 |
| Phospho DIA (LINCS P100 Plate 34) | 43.1 GB | 7,260 | 7,260 | 96 | 4 | 50 | 3h 33m | $15.01 | $0.00207 | 1.76 |
| | | | | | 10 | 10 | 4h 51m | $11.18 | $0.00154 | 2.40 |
| | | | | | 10 | 20 | 3h 12m | $12.21 | $0.00168 | 1.59 |
| | | | | | 10 | 30 | 2h 24m | $9.08 | $0.00125 | 1.19 |
| | | | | | 25 | 1 | 9h 53m | $10.34 | $0.00142 | 4.90 |
| Triple-proteome Pyrococcus furiosus proteome | 39.1 GB | 46,676 | 81,428 | 16 | 10 | 30 | 6h 5m | $38.43 | $0.00047 | 0.27 |
| Triple-proteome | 67GB | 100,360 | 111,222 | 16 | 10 | 50 | 7h 28m | $78.55 | $0.00071 | 0.24 |

**Supplementary table 4: Run time and cost for the four datasets presented in this study.** The dataset size represents the file size of the CSV file exported from Skyline and containing the chromatogram information for all peptides. The number of peptides, precursors and replicates contained in the dataset is shown. The analysis was run with several values for the number of VM cores and the number of CPUs per core. The corresponding cost and run time is provided.

| Category | | Location | Description |
|---|---|---|---|
| Avant-garde Terra implementation Release v1.0 | | https://app.terra.bio/#workspaces/lincs-phosphodia/Avant-garde_Production_v1_0 | Workspace that includes the methods to run the AvG algorithm. |
| Code | Avant-garde Terra implementation Release v1.0 | https://github.com/broadinstitute/Avant-garde-Terra | Github repository for the Terra workflow with code, documentation and description of the Terra workflow. |
| | Avg_utils | https://github.com/SebVaca/avg_utils | Avant garde python package containing the underlying functions required for the workflow |
| Tutorial | Tutorial description | https://github.com/broadinstitute/Avant-garde-Terra/wiki/Tutorial | Step-by-step instructions to run the AvG Terra workflow. |
| | Tutorial workspace | https://app.terra.bio/#workspaces/lincs-phosphodia/Avant-garde_Tutorial | Terra workspace with tutorial step-by-step instructions, data, and results. |
| Documentation | Documentation for the Avant-garde Terra implementation | https://github.com/broadinstitute/Avant-garde-Terra/wiki | Documentation describing AvG's requirements, inputs and outputs. |
| Case studies | Triple-proteome | https://app.terra.bio/#workspaces/lincs-phosphodia/Avant-garde_TripleProteome | |
| | Triple-proteome searched with Pyrococcus furiosus proteome | https://app.terra.bio/#workspaces/lincs-phosphodia/Avant-garde_TripleProteome_Pyrococcus_furiosus | |
| | LINCS phospho-enriched PC3 dataset | https://app.terra.bio/#workspaces/lincs-phosphodia/Avant-garde_LINCS_PC3 | |

**Supplementary table 5: Resources URLs for the Avant-garde Terra implementation release v1.0 workspace, code, tutorial and documentation.**

### Supplementary Information

#### Principles of the Avant-garde Terra framework

AvG algorithm was adapted to the Terra cloud-based infrastructure (http://app.terra.bio), a user-friendly Google Cloud-based platform for large-scale data analysis and sharing, as an accessible and standardized resource to the wider community. The new AvG workflow on Terra is optimized for large DIA datasets and fully automates all processing steps so that no user interaction is required after uploading the data to be analyzed. Terra takes care of the workflow management and facilitates the build-up of long-running pipelines that comprise a large number of tasks and take days or weeks to complete. To do this, each step of the AvG algorithm was implemented as a task, and the workflow represents the series of tasks (a pipeline) where outputs of a given task are the inputs of the next task. The workflow is written in the Workflow Description Language (WDL) and executes dockerized tools in R and python for pre-processing, running the AvG algorithm, and post-processing, all of which are available in a public docker image (Figure 1 and Supplementary Figure 1).

The workflow takes a CSV file containing chromatogram signal information exported from Skyline (Supplementary Figure 1). The AvG workflow converts the input file into an Apache Parquet file format that makes it compatible with a cloud-based data analysis platform. Parquet is an open-source file format designed for efficient and performant flat columnar storage; it is optimized to work with complex data in bulk and features different ways for efficient data compression, encoding, and partitioning. Taking advantage of Parquet's data partitioning features, the AvG workflow is highly parallelizable, and the data can be distributed across a user-defined number of nodes. Within each node, the AvG algorithm is run on each data partition individually. For further efficiency, a second parallelization occurs at this stage so that multiple CPUs on each node can be used to process several analytes at a time. The user can modulate efficiency by optionally defining the number of CPUs on each node as an input parameter. The final task of the workflow gathers the individual results from all analytes into output CSV files. These results can then be uploaded into Skyline to refine peak boundaries and filter out transitions unsuitable for quantification.

For users, the AvG Terra workspace contains all that is needed to run the algorithm. This includes the AvG workflow, the user-defined analysis parameters, and the results. Additionally, reproducible results can be ensured by Terra's version control system that accounts for the input dataset, algorithm, parameters, and results. The cloud-based AvG workflow harnesses the computational power of cloud-based computing to provide reproducible, user-friendly, and efficient processing of large DIA datasets with greater accessibility than our previous workflow. Its distribution on Terra allows anyone, regardless of coding experience, to perform efficient AvG analysis for large DIA datasets without the need to implement the source code locally or to access a large-scale computing resource.

#### Evaluation of the Terra implementation of Avant-garde

We first evaluated the Terra implantation of Avant-garde using a spike-in dataset. For this 95 synthetic peptides at known individual concentrations were spiked into a HEK293T digest to generate a five-point calibration curve. Each sample contained 1 µg HEK293T digest and 6.75, 13.5, 27, 54, or 108 ng total amount of heavy peptides. The five samples were analyzed in DIA mode in triplicates. In total, 15 runs were analyzed. The data was loaded into Skyline and the data was curated using the Skyline External Tool version and the Terra Workflow version of AvG. The relative quantification of the two datasets matched (Supplementary figure 2) showing that the results between the two implementations of AvG provide the same results for the relative quantification.

We applied the AvG Terra workflow to a triple-proteome sample consisting of four mixture samples of three complex proteomes (Human, E. Coli and Yeast) that was described in the original AvG paper[1]. Briefly, the total amount of protein and the proportion of the human proteome were kept constant in all samples, while the proportion of E. coli and yeast varied. Pairwise combinations of the samples result in 'ground truth' ratios ranging from 1.2-fold to 10-fold, plus a constant 1:1 ratio of human peptides for all possible comparisons. This dataset enabled the evaluation of the reproducibility and accuracy across many MS runs having different sample compositions, with some compositions more prone to interference than others. Additionally this dataset has a large dynamic range of peptide abundance and mimics ratios close to typical thresholds of biological significance for the evaluation of DIA analysis tools. Refinement with AvG provided high quantification accuracy and precision. We evaluated the relative quantification accuracy, by choosing sample C as the denominator, and we calculated the mean area for each run (A,B or D) to the mean area of sample C. We are expecting to have 3 pairwise comparisons producing three expected ratios for the yeast proteome (-8, -2, 1.2), three expected ratios for the E. Coli proteome (-3, 2.3 and 3.3) and one for the Human proteome (1). The results of this evaluation show excellent accuracy, as the measured ratios matched very closely the expected ratios (Supplementary Figure 3, Supplementary Figure 4 and supplementary Table 1). Overall, in the entire dataset we obtain a mean and median absolute percent error equal to 17.0% and 10.3%, which indicates that the measured ratios are very close to the expected values. Similarly, the mean and median coefficient of determination were respectively $r^2$=0.97 and $r^2$=1. Additionally, the distribution of the percent error for each individual pairwise comparison  shows similar values for the accuracy  indicating that we obtain excellent accuracy across all samples measured (Supplementary Figure 4). Furthermore, we were able to validate more than 34,000 precursors in the whole dataset (Supplementary Table 1 and 2). For the entire dataset we obtain excellent reproducibility with a mean and median CV of 15.3% and 12.3% respectively (Supplementary Table 2). Additionally, the distribution of CVs for each individual sample analyzed in quadruplicates also showed excellent reproducibility indicating that we obtain excellent reproducibility for all MS runs (Supplementary Figure 5 and Supplementary table 2).

To further evaluate the confidence level of the results, we searched the Triple-species proteome samples with a Pyrococcus furiosus proteome strectral library, which provides a method for detecting random hits [2]. No Pyrococcus furiosus proteins were present in the sample and we did this to verify that we see a negative result when searching the wrong library. The spectral library for the Pyrococcus furiosus peptides was predicted using a deep neural network tool, Prosit[3], that provided chromatographic retention time and fragment ion intensity predictions. As expected, the score distribution of the Pyrococcus furiosus and the decoy peptides overlapped (Supplementary figure 6) and of the 497,263 DIA features, corresponding to 21074 peptides, only 15 Pyrococcus furiosus DIA features were cumulatively identified, corresponding to 10 peptides, yielding an 'intrinsic' false discovery rate of 0.003% (Supplementary Figure 6). Additionally, zero Pyrococcus furiosus peptides were validated when using a posterior error probability lower than 0.05, resulting in a 'intrinsic' false discovery rate of 0% .

Finally, a run time and cost overview of running the AvG Terra workflow on four datasets can be found in the Supplementary Table 4. The running time and cost of the analysis depend on several parameters. Focusing on the LINCS phospho-enriched PC3 dataset, we have discovered that the most influencing parameters are the size of the input file, the number of analytes, the number of virtual machine cores, the number of CPUs per core (Supplementary table 4). Additionally, reducing the size of the time window of the extracted chromatograms highly reduces the cost and time. Overall in this study, the cost of running this pipeline ranged between $0.00047 and $0.00207 per peptide and took between an effective processing time between 0.24 and 11.88 seconds per peptide.

### Supplementary Methods

Detailed methods for the Avant-garde algorithm can be found in the original paper[1].

Spike-in calibration curve of heavy peptide dataset

The 96 peptide dataset was described in the original AvG paper[1]. Briefly, a mixture containing 95 synthetic peptides at known individual concentrations was spiked into a HEK293T digest to generate a five-point calibration curve. Each sample contained 1 µg HEK293T digest and 6.75, 13.5, 27, 54, or 108 ng total amount of heavy peptides. The solution of heavy peptides used for this experiment was a mixture of 95 peptides that were combined in different concentrations to get a concentration-balanced mixture. Given that not all of these peptides have the same response factor, we adjusted the concentration of each peptide to ensure its detection. The five samples were analyzed in DIA mode in triplicates. In total, 15 runs were analyzed. The data was loaded into Skyline and the data was curated using the Skyline External Tool version and the Terra Workflow version of AvG. Skyline was set with the following parameters: precursor charges, 2, 3, 4; ion charges, 1, 2; ion types, y; ion match tolerance, 0.5 m/z; product mass analyzer, Orbitrap; MS1 resolving power, 60,000 at 200 m/z; MS2 resolving power, 30,000 at 200 m/z: chromatograms were extracted 10 min each side of the predicted retention time. The 25 most intense transitions for each peptide were extracted and the peak boundaries were determined by Skyline. The data was filtered to keep only DIA features having an AvG score higher than 0.1.

Triple-proteome dataset

A spectral library was created from DDA MS runs. The DDA data were interpreted using the Spectrum Mill software package v6.0 Pre-release (Agilent Technologies, Santa Clara, CA, USA). MS/MS spectra were excluded from searching if they failed the quality filter by not having a sequence tag length>0 (that is, minimum of two masses separated by the in-chain mass of amino acid) or did not have a precursor MH+ in the range of 400–4,000. MS/MS spectra were searched against a database consisting of a human Uniprot database containing a set of 150 common laboratory contaminant proteins (UniProt.human.20141017.RNFISnr.150contams). The scoring parameters were ESI-QEXACTIVE-HCD-v2. Mass tolerance was fixed at ±10 p.p.m. and ±20 p.p.m. for precursor and product ions respectively. The minimum matched peak intensity was 50%, and trypsin specificity with up to 2 missed cleavages. Fixed modifications were carbamidomethylation at cysteine. Variable modifications were acetylation of protein N-termini, oxidized methionine, phosphorylation (S, T, Y) with a precursor MH+ shift range of −18 to 250 Da. Spectrum Mill autovalidation module was used to achieve an FDR <1%. The results were exported as a pep.xml file and imported into Skyline to generate the spectral library. The DIA data was analyzed using Skyline. Raw files were converted to MzML and demultiplexed using MSConvert. Chromatograms were extracted using Skyline with the following parameters: precursor charges, 2, 3, 4; ion charges, 1, 2; ion types, y, b; ion match tolerance, 0.5 m/z; product mass analyzer, centroided; tandem mass spectrometry mass accuracy, 10 ppm: chromatograms were extracted 5 min each side of the predicted retention time. The ten most intense transitions for each peptide were extracted and the peak boundaries were determined by Skyline, using a set of 15 retention time standard peptides for retention time prediction. The data were refined a posteriori by AvG to select at least four transitions per peptide and to correct the peak boundaries. The data were scored and filtered to less than 1% FDR using Percolator v 3.0 (the SLS score , mass error score, PSS score, MPRA score were used as features for Percolator). The data was filtered at a q-value<0.01. The percent error was calculated as the absolute value of the difference between the measured and the expected ratio divided by the expected ratio.

Pyroccosus Furiosus dataset

The Pyroccosus Furiosus reviewed protein database was downloaded from Uniprot (Proteome ID: UP000001013, reviewed: yes, download date: 2021/04). The Fasta file was used as a background proteome in Skyline version Skyline-Daily (64-bit) 21.0.9.139. The peptides in silico digested were selected with a Trypsin enzyme, maximum missed cleavages of 2, peptide length between 8 and 25, Methionine-containing peptides were excluded. For all the peptides we created a spectral library where spectra were predicted using the Prosit implementation in Skyline. An NCE of 27 was used and an iRT calculator was aligned using 12 human peptides that were verified to be present in the samples in high-abundance. Skyline was set with the following parameters: precursor charges, 2, 3, 4; ion charges, 1, 2; ion types, y, b; ion match tolerance, 0.5 m/z; product mass analyzer, centroided; tandem mass spectrometry mass accuracy, 10 ppm: chromatograms were extracted 5 min each side of the predicted retention time. The ten most intense transitions for each peptide were extracted and the peak boundaries were determined by Skyline, using a set of 12 retention time standard peptides for retention time prediction. The data were refined using the AvG Terra Workflow. The data was validated using Percolator v 3.0 (the SLS score , mass error score, PSS score, MPRA score were used as features for Percolator) at a q.value < 0.01 at the PSM level (in this case DIA features) or using the posterior error probability <0.05 (PEP<0.05).

LINCS phospho-enriched PC3 dataset.

A phospho-enriched sample dataset was used to evaluate the Terra AvG framework. Detailed sample preparation protocols and nanoLC-MS/MS methods can be found online at https://panoramaweb.org/wiki/LINCS/Overview%20Information/page.view?name=sops and in our previous publications [4,5]. Briefly, PC3 were cultured and treated as described in detail in our previous study. PC3 cells were cultured in RPMI 1640 medium containing 1 mM sodium pyruvate and 10 mM HEPES (Thermo Fisher Scientific). Cells were plated onto six-well plates for 24 hours, expanded to near confluence, and treated by adding the drug of interest diluted media at the desired concentration. For cell harvest, lysis buffer (8 M Urea, 75 mM NaCl, 50 mM Tris HCl, pH 8.0, 1 mM EDTA, 2 µg/ml aprotinin, 10 µg/ml leupeptin, 1 mM PMSF, 10 mM NaF, Phosphatase Inhibitor Mixture 2 and Phosphatase Inhibitor Mixture 3) was added in each well and cells were collected via scraping. Samples were lysed for 15 minutes at room temperature and then vortexed, followed by an additional 15-minute incubation before freezing. Upon thawing, lysates were centrifuged at 15,000 x *g,* 15°C for 15 minutes to pellet cell debris and extract protein slurry. Protein concentration was measured using the 660 protein assay (Pierce, 22660). All samples (~500 ug each) were normalized to a protein concentration of 1.25 µg/µl. Proteins were reduced, alkylated, and digested overnight with sequencing-grade modified trypsin at an enzyme:substrate ratio of 1:50 (Promega, V511X, Madison, WI). Upon quenching, samples were desalted using reversed-phase SPE (Waters, 186002319). Peptides were eluted with 50%ACN/0.1%TFA and lyophilized. Peptides were reconstituted in 80%ACN/0.1%TFA. Phosphopeptides were enriched using $Fe^{3+}$ IMAC cartridges (AssayMAP Bravo, Agilent, Santa Clara, CA) following standard protocol. Phosphopeptides were desalted using AssayMAP Reverse Phase cartridges (Agilent, G5496-60033) and lyophilized. Before MS analysis, a set of synthetic isotope-labeled peptides were spiked into the samples to allow for quantitation. Chromatograms were extracted using Skyline. Skyline was set with the following parameters: precursor charges, 2, 3, 4; ion charges, 1, 2,3; ion types, y, b; ion match tolerance, 0.5 m/z; product mass analyzer, centroided; tandem mass spectrometry mass accuracy, 10 ppm: chromatograms were extracted 5 min each side of the predicted retention time. The ten most intense transitions for each peptide were extracted and the peak boundaries were determined by Skyline. The data were refined using the Terra implementation of Avant-garde.

<u>LC-MS/MS</u>

The samples were analyzed using a Orbitrap Q-Exactive HF Plus (Thermo Fisher Scientific) mass spectrometer coupled to a nanoflow Proxeon EASY-nLC 1000 UHPLC system (Thermo Fisher Scientific). The mass spectrometer was used in positive mode and was equipped with a nanoflow ionization source (James A. Hill Instrument Services); the spray voltage was set at 2.00 kV. The liquid chromatography system, the column, and the electrospray voltage source (platinum wire) were connected using a stainless steel cross (360 µm, IDEX Health & Science, UH-906x). The column was heated to 50 °C. A volume of 3 µl was injected onto an in-house packed 20 cm × 75 µm diameter C18 silica picofrit capillary column (1.9-µm ReproSil-Pur C18-AQ beads, Dr Maisch, r119.aq; Picofrit 10-µm tip opening, New Objective, PF360-75-10-N-5). The mobile phase had a flow rate of 250 nl min−1 and consisted of 3% ACN/0.1% FA (solvent A) and 90% ACN/0.1% FA (solvent B). The column was conditioned before each sample injection. Peptides were separated using the following liquid chromatography gradient: 0–3% B in 3 min, 5–40% B in 50 min, 40–90% B in 1 min, stay at 90% B for 5.5 min, and 90–50% B in 30 s. DDA and DIA data were acquired on the same instrument. For the MS1 scans, the resolution was set at 60,000 at 200 m/z and the automatic gain control (AGC) target was 3 × 106 with a maximum inject fill time of 20 ms. For DDA, MS2 scans on the top 12 peaks doubly charged and above were acquired at a resolution of 15,000, and an AGC target of 5 × 104 with maximum inject fill time of 50 ms. Isolation widths were set to 1.5 m/z with a 0.3 m/z offset. The normalized collision energy (NCE) was set to 27 and dynamic exclusion was set to 10 s. For DIA, an overlap DIA method was used with 56 × 22 m/z isolation windows covering the 400–1,000 m/z range. In this method, the isolation windows in two consecutive cycles have an offset of 11 m/z. The default charge state was 4, the resolution was 30,000 at 200 m/z, the AGC target was 1 × 106, the maximum inject fill time was 50 ms, the loop count was 27 and the NCE was set to 27.

### Tutorial for running Avant-garde on Terra

Avant-garde is a tool for automated signal refinement of DIA and other targeted mass spectrometry data[1]. Avant-garde can be used alongside existing tools for peptide detection to refine chromatographic peak traces of peptides identified in DIA data by these tools. Avant-garde ensures their confident and accurate quantitation by removing interfering transitions, adjusting integration boundaries, and scoring peaks to control the false discovery rate. The improved Avant-garde workflow described here is deployed on Terra (https://app.terra.bio/), a Google cloud-based platform for large-scale data analysis and sharing, as a fully-automated, reproducible, scalable, and user-friendly workflow.

This tutorial provides step-by-step guidance on how to set up, run, and obtain results from the Avant-garde workflow on Terra. The example data used for this tutorial is the small reduced-representation phosphoproteomic DIA dataset of 96 phosphopeptides x 15 samples described in Vaca Jacome et al. 2020[1]. The input dataset is called `Tutorial_AvantGardeDIA_Export.csv` and it can be extracted from the zip file called `Tutorial_AvantGardeDIA_Export.zip`(https://github.com/broadinstitute/Avant-garde-Terra/blob/master/tutorial/Tutorial_AvantGardeDIA_Export.zip) found in the `tutorial` subdirectory of the Avant-garde-Terra Github repository.

The Terra workspace created by this tutorial is called Avant-garde_Tutorial (https://app.terra.bio/#workspaces/lincs-phosphodia/Avant-garde_Tutorial).

#### 1. Requirements

1. You need to sign up for a Terra account, if you don't have one already. Signing up for a Terra account is free and only requires a Google account (which is also free to sign up for). Instructions for signing up for a Terra account can be found here: https://support.terra.bio/hc/en-us/articles/360034677651-Account-setup-and-exploring-Terra.
2. You will then need to create a Google Cloud Platform (GCP) Billing Account and link it to Terra to create a Terra Billing Project; step-by-step instructions for doing so can be found here: https://support.terra.bio/hc/en-us/articles/360026182251-How-to-set-up-billing-in-Terra. New users to GCP billing are eligible to claim $300 in free GCP credits for exploring Terra; see here: https://support.terra.bio/hc/en-us/articles/360046295092 for more details.

#### 2. Clone the Avant-garde production workspace on Terra

Clone the Avant-garde production workspace (https://app.terra.bio/#workspaces/lincs-phosphodia/Avant-garde_Production_v1_0) into your Terra account.

1. Navigate to the Avant-garde_Production_v1_0 workspace (https://app.terra.bio/#workspaces/lincs-phosphodia/Avant-garde_Production_v1_0). Click on the 3 vertical dots in the top right and then click "Clone":

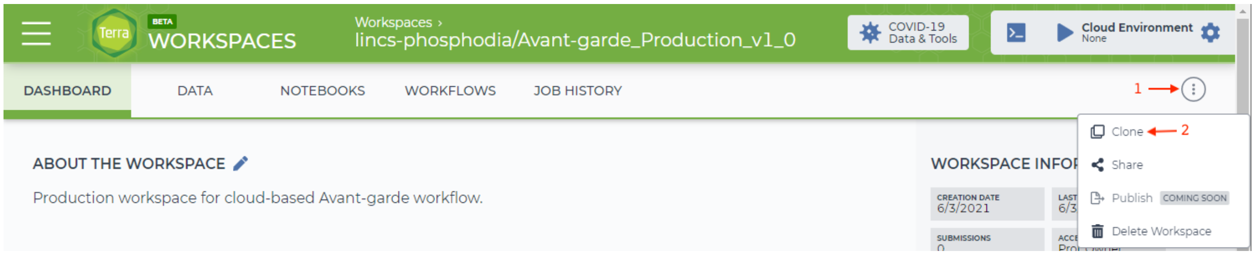


Tutorial figure 1: Cloning the Avant-garde production workspace on Terra

2. Name your new workspace and assign it to your Terra billing project, and then click "CLONE WORKSPACE":

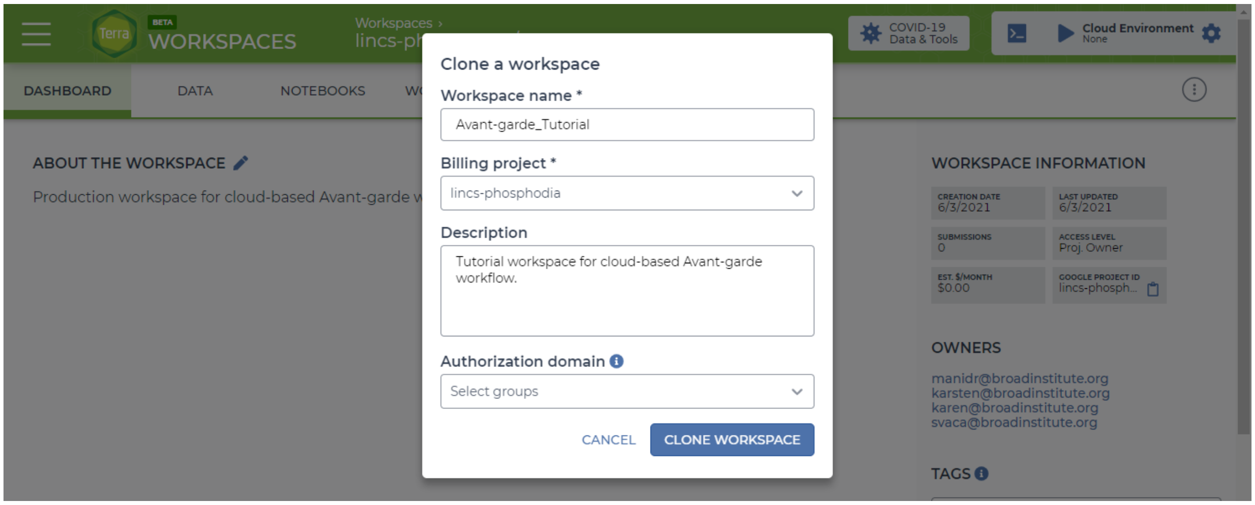


Tutorial figure 2: naming your new Avant-garde workspace

3. Upload data and parameters file to the workspace's Google bucket

Navigate to the DATA tab of your new workspace and click on "Files" on the left menu under OTHER DATA. Click on the blue "+" at the bottom right and upload the following two required inputs:

1. Avant-garde parameters file. Download `AvG_Params.R` from the Avant-garde-Terra Github repository in the templates (https://github.com/broadinstitute/Avant-garde-Terra/tree/master/templates) subdirectory, or from the DATA tab (https://app.terra.bio/#workspaces/lincs-phosphodia/Avant-garde_Production_v1_0/data) (under Files) of the Avant-garde production workspace. If required for your data, make any edits to the default parameters (most users will be able to use all defaults; see

Parameters (https://github.com/broadinstitute/Avant-garde-Terra/wiki/Parameters) documentation for more information). Upload `AvG_Params.R` to the DATA tab of your new workspace as described above.

2. CSV file containing chromatogram data. This can be generated as a report in Skyline using the template included in the Skyline installation of Avant-garde or here https://github.com/SebVaca/Avant_garde/blob/master/skyline_external_tool/AvG_skyline externaltool/tool-inf/AvantGardeDIA_Export.skyr. For this tutorial, the input CSV file used (`Tutorial_AvantGardeDIA_Export.csv`) can be found in the Avant-garde-Terra Github repository in the tutorial subdirectory (https://github.com/broadinstitute/Avant-garde-Terra/tree/master/tutorial) - it must first be extracted from the `Tutorial_AvantGardeDIA_Export.zip` archive.

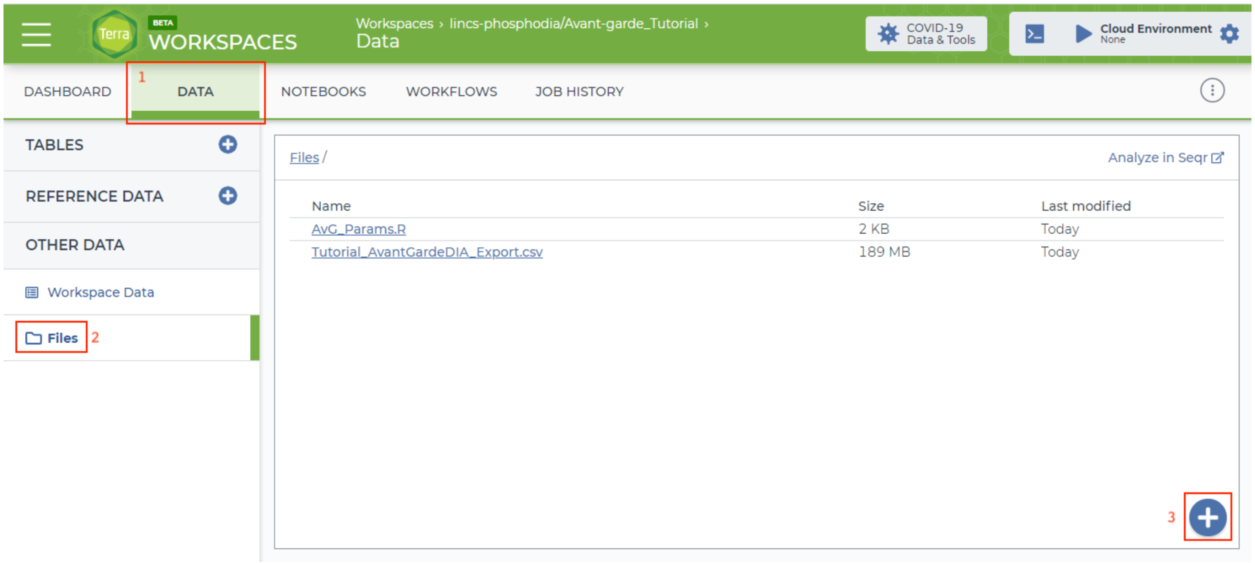


Tutorial figure 3: uploading data to the Terra workspace under the DATA tab

NOTE: If you already have either of these files stored in an existing Google Bucket on the GCP billing account linked to your Terra account, you can skip this step and simply copy and paste the gsutil URI for the given file into the appropriate input field shown in the next step.

#### 4. Run the Avant-garde workflow

Navigate to the WORKFLOWS tab and click on the "Avant-garde" workflow:

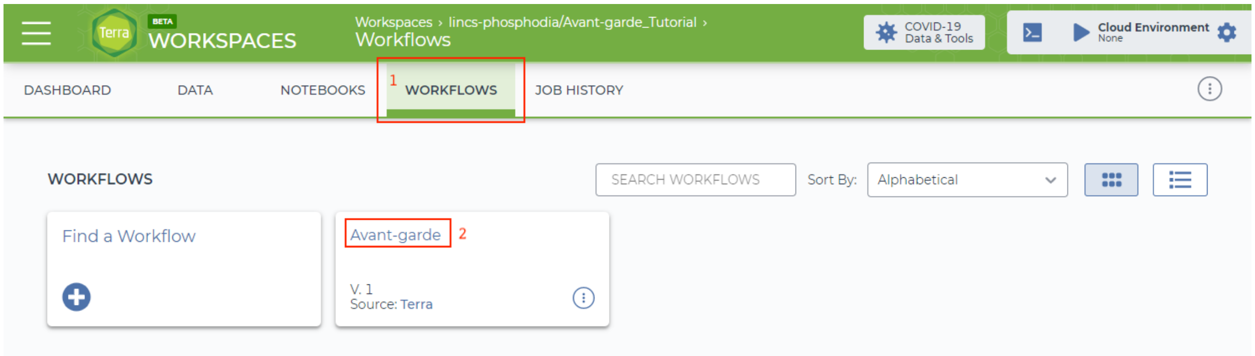


Tutorial figure 4: Navigating to the Avant-garde workflow

1. **IMPORTANT**: select the option to "Run workflow with inputs defined by file paths" at the top of the page.

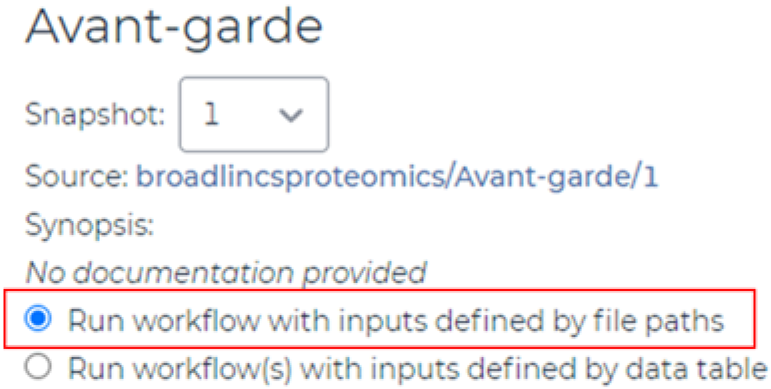


Tutorial figure 5: select the option to "Run workflow with inputs defined by file paths".

2. Fill in 6 required inputs. See `Avantgarde` workflow documentation(https://github.com/broadinstitute/Avant-garde-Terra/wiki/Avantgarde) for more details on the required inputs. For this small tutorial dataset, a chunk size of 4000 and 6 virtual machines for parallelization are chosen. For the `inputcsv` and `params_file` inputs, click on the folder icon labeled "Browse bucket files" to select `Tutorial_AvantGardeDIA_Export.csv` and `AvG_Params.R`, respectively, which were uploaded to the Google bucket in Step 3. Alternatively, if your file is stored in a different Google bucket, copy and paste the gsutil URI for that file into the respective input field (make sure it is in quotation marks).
3. Fill in optional inputs as necessary. See `Avantgarde` workflow documentation(https://github.com/broadinstitute/Avant-garde-Terra/wiki/Avantgarde) for more details on the optional inputs. For this small tutorial dataset, the default of 10 CPU is not necessary, so 4 CPU is written in the `num_cpu` field to override the default.
4. Click "SAVE" to save your inputs

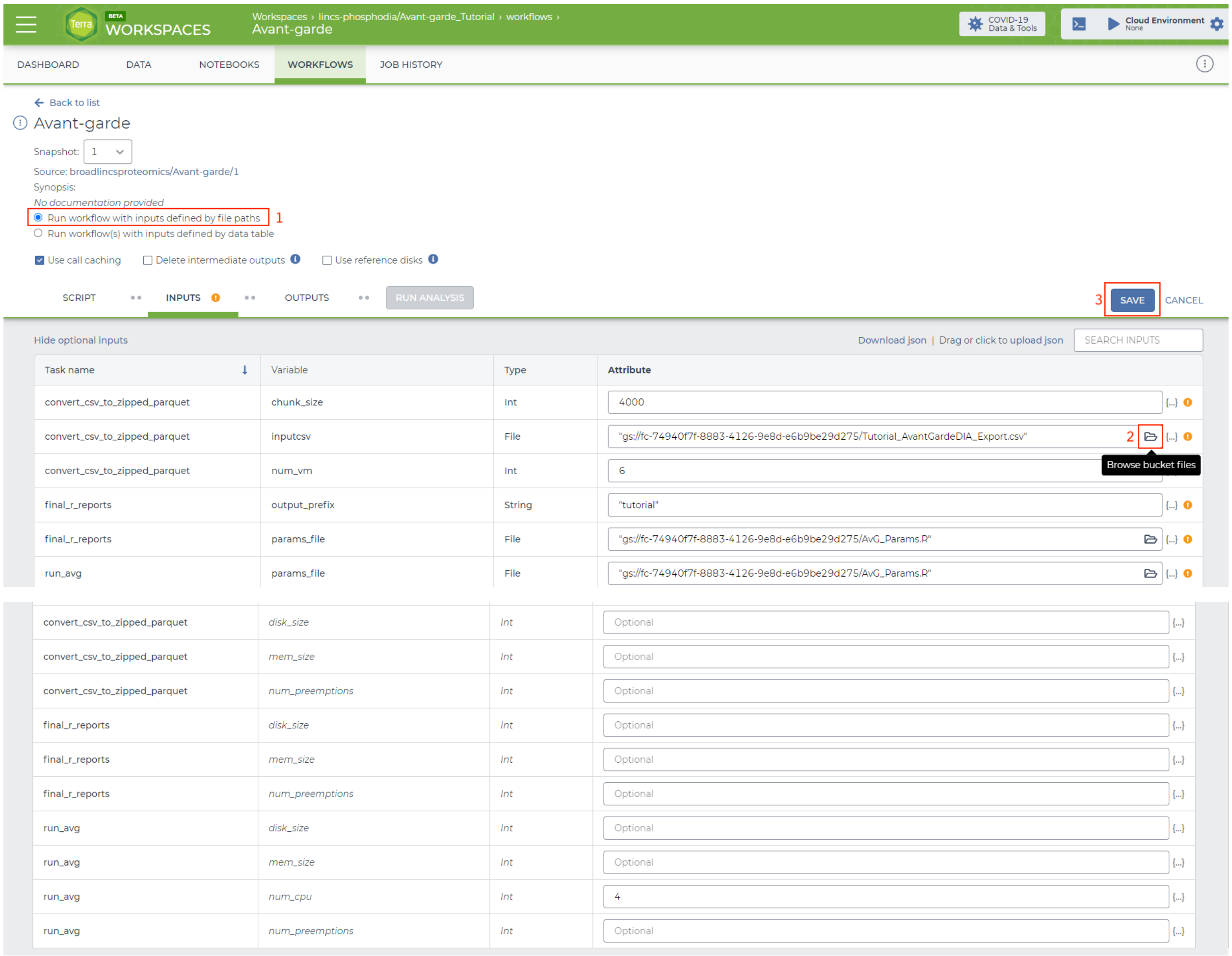


Tutorial figure 6: filling in required and optional inputs to the workflow

5. Click "RUN ANALYSIS" and then click "LAUNCH" when asked to confirm launch.

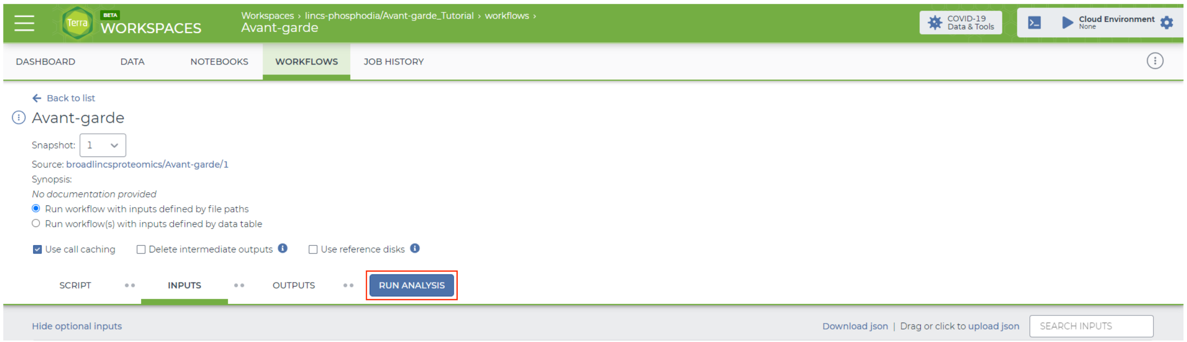


Tutorial figure 7: Click RUN ANALYSIS

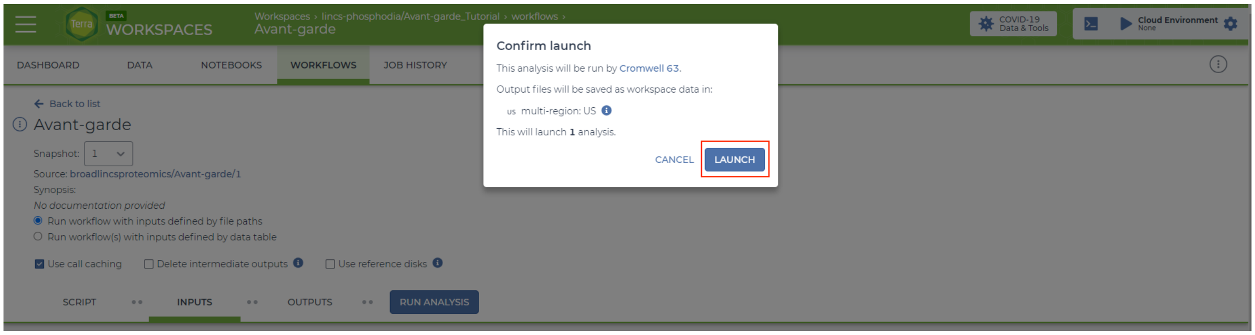


Tutorial figure 8: Click LAUNCH

6. You will be automatically redirected to the job submission page where you can monitor the analysis in real time. If you navigate away, you can return to the submission page by clicking on the JOB HISTORY tab.

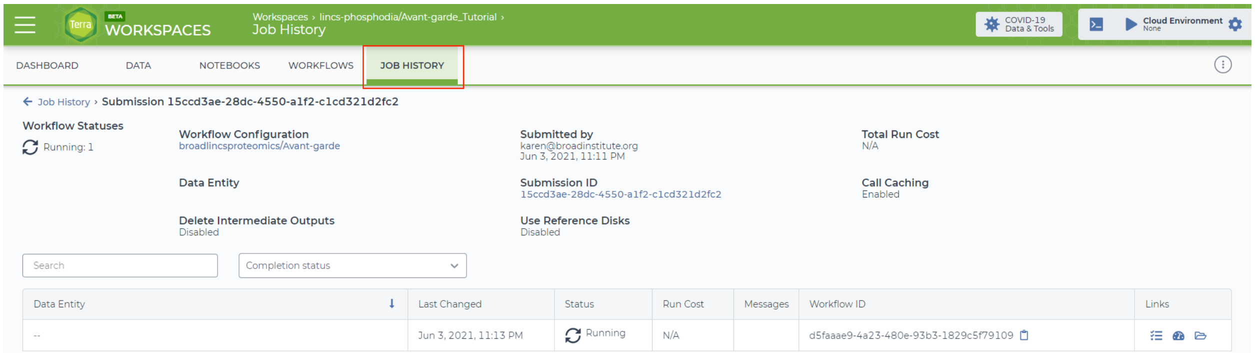


Tutorial figure 9: submission page for this run under the JOB HISTORY tab

#### 5. Downloading results

When the analysis is completed successfully, a green check mark will appear on the job submission page in the JOB HISTORY tab. Click on the Job Manager icon on the right to view details about the analysis, including results.

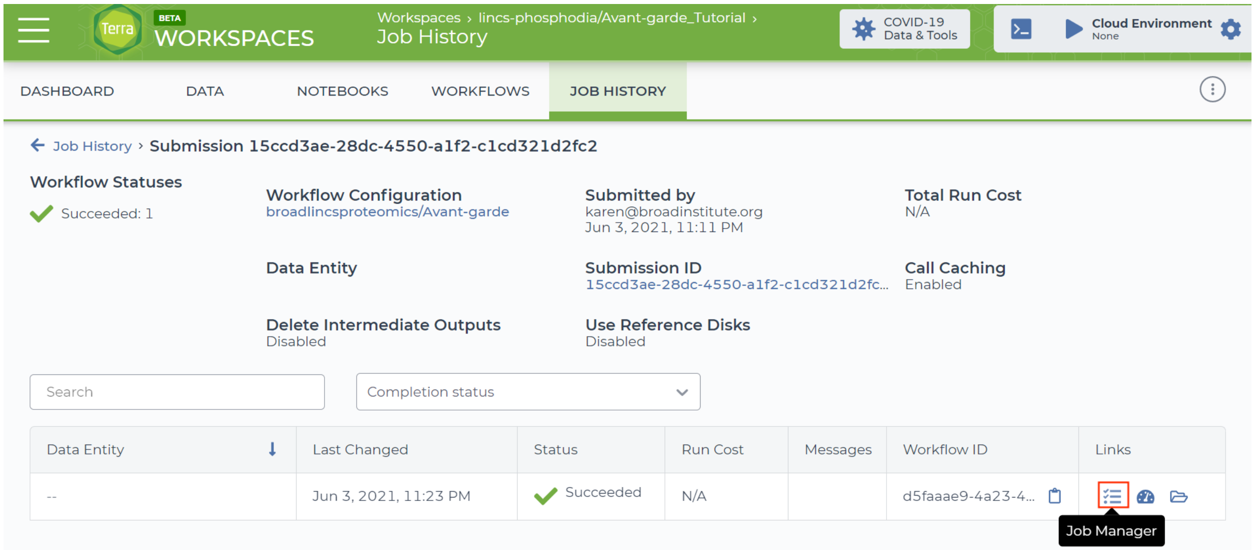


Tutorial figure 10: Successful job completion and Job Manager icon

The Job Manager page displays the inputs and outputs of all 3 tasks in the workflow. The final Avant-garde results are the output of the final task in the workflow, `final_r_reports`. Click on the icon under "Outputs" for the `final_r_reports` task and a pop-up will open showing the Google Bucket link to the output zip file containing the Avant-garde results. Click on this link to download the zip file containing the Avant-garde results.

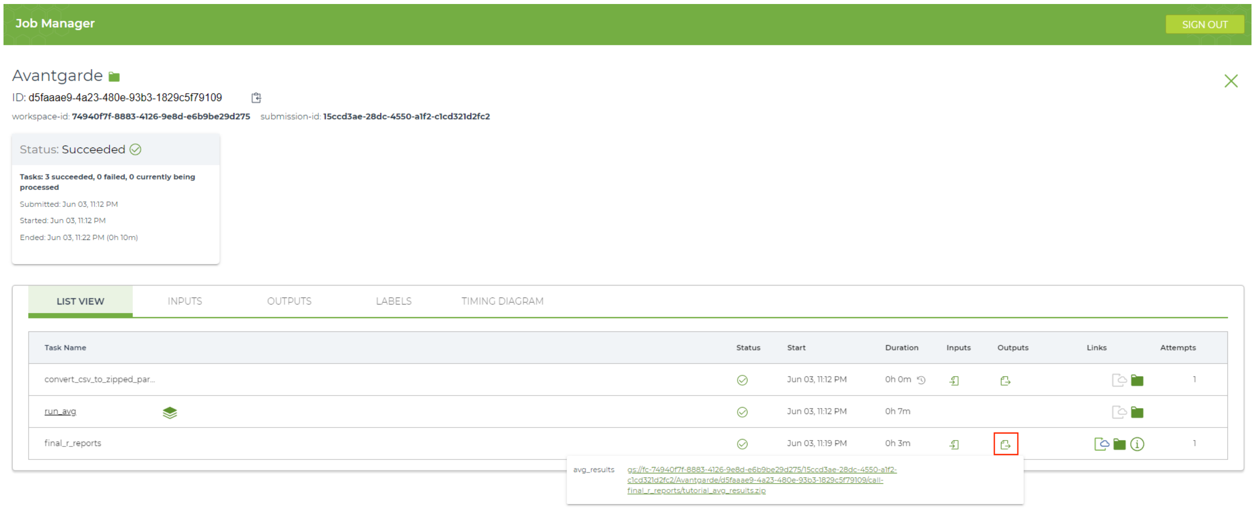


Tutorial figure 11: Locating Avant-garde results on Job Manager page

### References


1. Vaca Jacome, A. S. *et al.* Avant-garde: an automated data-driven DIA data curation tool. *Nat. Methods* **17**, 1237–1244 (2020).
2. Vaudel, M. *et al.* A complex standard for protein identification, designed by evolution. *J. Proteome Res.* **11**, 5065–5071 (2012).
3. Gessulat, S. *et al.* Prosit: proteome-wide prediction of peptide tandem mass spectra by deep learning. *Nat. Methods* **16**, 509–518 (2019).
4. Abelin, J. G., Patel, J., Lu, X., Feeney, C. M. & Fagbami, L. Reduced-representation phosphosignatures measured by quantitative targeted MS capture cellular states and enable large-scale comparison of drug-induced …. *Molecular & Cellular* (2016).
5. Litichevskiy, L. *et al.* A Library of Phosphoproteomic and Chromatin Signatures for Characterizing Cellular Responses to Drug Perturbations. *Cell Syst* **6**, 424–443.e7 (2018).
